# Structural Determinants and Biochemical Characterization of LORELEI as a GPI-Anchored Protein

**DOI:** 10.1101/2025.06.10.659000

**Authors:** Yanbing Wang, Xunliang Liu, Nicholas Desnoyer, Gregory Howard, Ravishankar Palanivelu

## Abstract

Glycosylphosphatidylinositol-anchored proteins play critical roles in plant development, reproduction, and environmental responses. However, their biochemical characterization in plants remains limited. LORELEI, a putative glycosylphosphatidylinositol-anchored protein involved in pollen tube reception and early seed development in *Arabidopsis thaliana*, has lacked direct biochemical evidence confirming its glycosylphosphatidylinositol anchoring. This study employed biochemical approaches to validate LORELEI as a glycosylphosphatidylinositol-anchored protein. We demonstrate that ectopic expression of wild-type LORELEI fused to a reporter protein in vegetative tissues associates with a detergent-resistant membrane fraction and is sensitive to glycosylphosphatidylinositol-specific cleavage enzymes, thereby confirming that it is a glycosylphosphatidylinositol-anchored protein. Moreover, we show that mutations in the ω-sites or the glycosylphosphatidylinositol attachment signal of LORELEI disrupt its membrane localization, highlighting the necessity of these structural elements for proper glycosylphosphatidylinositol anchoring. Analysis of a reporter-fused LORELEI protein in the loss of function mutant of *glycosylphosphatidylinositol 8*, which encodes the catalytic subunit of glycosylphosphatidylinositol transamidase complex involved in glycosylphosphatidylinositol anchor addition, results in the appearance of prominent higher molecular weight bands, further supporting its role in glycosylphosphatidylinositol anchoring of the LORELEI protein. This study provides direct biochemical evidence confirming LORELEI as a glycosylphosphatidylinositol-anchored protein and sheds light on the structural determinants required for its glycosylphosphatidylinositol anchoring. Additionally, our study demonstrates that heterologous ectopic expression of LORELEI in vegetative tissues provides a viable alternative for the biochemical characterization of glycosylphosphatidylinositol-anchored proteins predominantly expressed in hard-to-access and small-sized female gametophytes. Our findings underscore the role of GPI anchoring in membrane localization and biosynthesis of GPI-anchored proteins in plants.

## Introduction

In *Arabidopsis thaliana*, LORELEI (LRE) is a putative GPI-anchored protein (GPI-AP) expressed in the female gametophyte before fertilization and in the developing seed after fertilization. Genetic studies show that LRE is essential for fertilization—loss-of-function mutations cause severe pollen tube reception defects, reducing seed yield by ∼80% and delaying seed development (Capron et al. 2008; Tsukamoto et al. 2010; Liu et al. 2016; Wang et al. 2017). LRE belongs to a small family of four LORELEI-LIKE GPI-anchored proteins (LLGs) involved in reproduction, growth, immunity, and stress responses in Arabidopsis (Li et al. 2015; Shen et al. 2017; Feng et al. 2018; Ge et al. 2019; Liu et al. 2021; Noble et al. 2022). In Arabidopsis, *LLG1* is broadly expressed, *LLG2* and *LLG3* expression is confined to pollen and pollen tubes, and LRE is expressed in female tissues (Li et al. 2015; Ge et al. 2019; Noble et al. 2022). These genes are evolutionarily conserved and have both core and species-specific functions across angiosperms (Noble et al. 2022).

GPI-APs are vital for fertility in flowering plants, supporting male and female gametophyte development (Desnoyer and Palanivelu, 2020). They also play roles in root and root hair development, shoot regeneration, tissue morphogenesis, seed coat formation, vascular growth, and stomatal regulation, likely by influencing cell division, polar expansion, cell wall synthesis, and signal transduction (Yeats, Bacic and Johnson, 2018; Zhou, 2019; Lin et al. 2022). Despite their importance, biochemical characterization of plant GPI-APs remains limited and is often inferred from studies in mammals, yeast, and protists (Desnoyer and Palanivelu, 2020).

In mammalian cells, GPI-AP biosynthesis starts with a preproprotein containing an N-terminal ER-targeting signal peptide and a C-terminal GPI attachment signal (GAS) (Kinoshita, 2020). The GPI anchor is added at the omega (ω) site, located just after the GAS sequence (Fig. 1). Once in the ER, the signal peptide is cleaved, forming a proprotein. GPI transamidase (GPI-T) then transfers the GPI moiety to the ω site, replacing the GAS and producing a mature GPI-AP (Kinoshita, Fujita and Maeda, 2008; Kinoshita, 2020). This involves cleavage by PIG-K, generating a thioester-linked intermediate, followed by transamidation via GPAA1, forming a stable amide bond with the GPI anchor (Kinoshita, 2020).

**Fig. 1.**
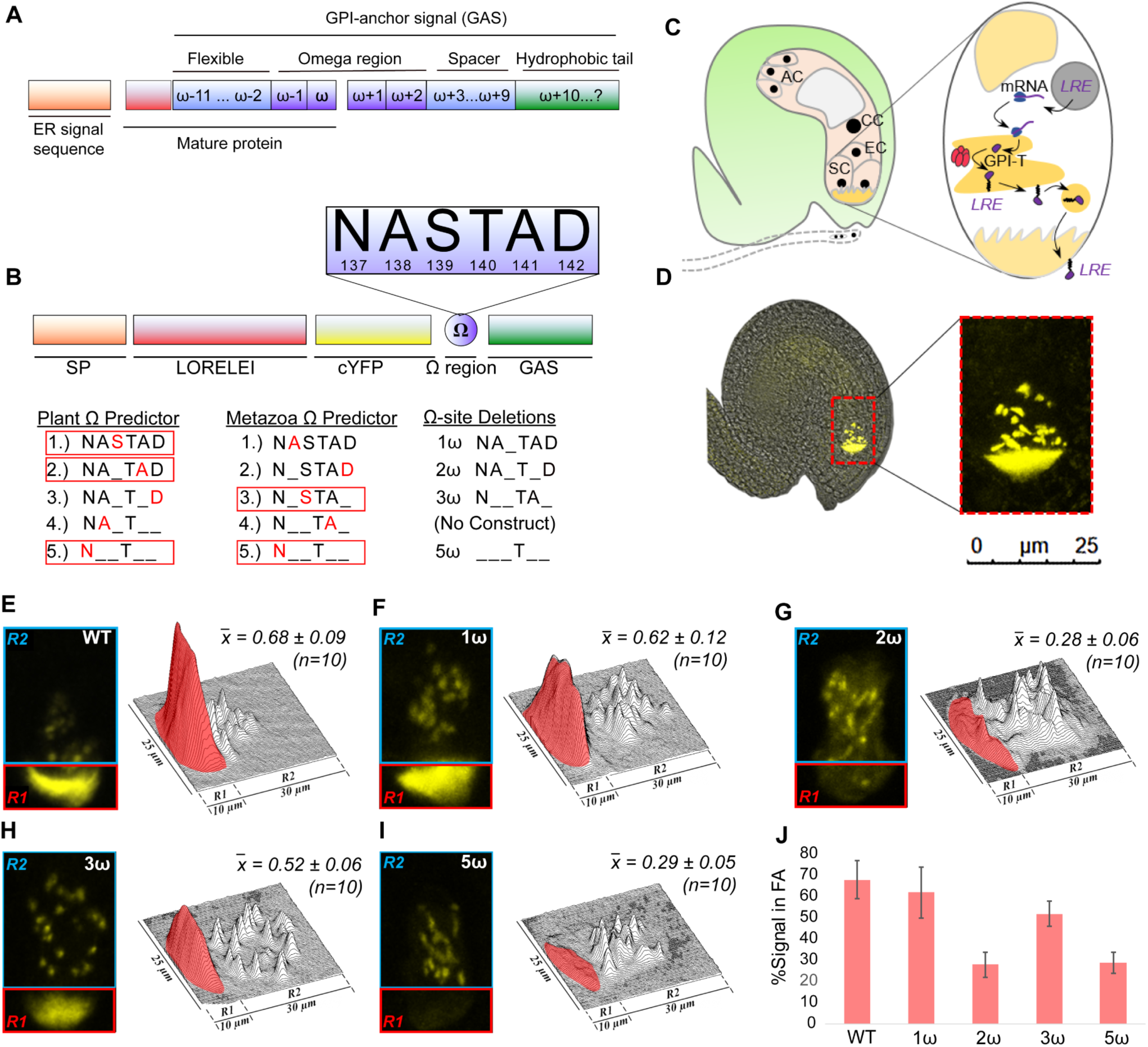
Deletion analysis of predicted GPI ω-Sites reveals Canonical and Cryptic ω Sites in LORELEI. A. Diagram detailing the key domains in LRE nascent protein produced in the ER, including the GPI anchor addition domains – ω site and the GPI Addition Sequence (GAS). B. The ω sites predicted by the two predictor sites and the various deletions of the ω sites are shown. C. The biosynthesis of LRE protein in the female gametophyte and the eventual localization of the LRE protein in the filiform apparatus of the synergid cells are shown. AC, antipodal cells (3); CC, central cell (1); EC, egg cell (1); SC, synergid cells (2). D. Left: A merged image (bright field and fluorescent) of an ovule expressing *pLRE:LRE-cYFP* is shown. Right: A close-up view of the portion of the ovule marked in red rectangle in the ovule image to the right. A majority of the cYFP signal is in the filiform apparatus within the syngergid cells and the rest of the cYFP signal is localized in unknown organelles in the cytoplasm of the synergid cells. E – I. Localization of *pLRE:LRE-cYFP* (E), *pLRE:LRE-cYFPΔ1ω* (F), *pLRE:LRE-cYFPΔ2ω* (G), *pLRE:LRE-cYFPΔ3ω* (H), and *pLRE:LRE-cYFPΔ5ω* (I) in the synergids of the female gametophyte. Left panels, representative image showing the localization of each fusion protein in the micropylar region of the ovules. R1 and R2, regions of interest 1 and 2, respectively, in the filiform apparatus (red box) and the cytoplasm of synergid cells (cyan box). Right panels, surface plots showing quantification of cYFP signal in the region of filiform apparatus (R1) compared to the total cYFP signal in the synergid cells (R1 + R2). n, number of ovules analyzed. Bar = 25 μm. J. Bar graph showing % signal intensity in the filiform apparatus (FA) of synergid cells of indicated constructs with or without predicted ω site(s) mutations.

In plants, GPI-AP biosynthesis likely follows a similar pathway (Ellis et al. 2010; Cheung et al. 2014). The GPI moiety assembles on the cytoplasmic ER face, flips into the lumen, and is transferred to the ω site by GPI-T after GAS cleavage. Following post-translational modifications, GPI-APs undergo lipid remodeling in the Golgi and are transported to membrane microdomains via vesicles (Ellis et al. 2010; Mañuel and Howard, 2016). In Arabidopsis, AtPGAP1-mediated remodeling is essential for efficient plasma membrane (PM) trafficking (Bernat-Silvestre et al. 2021). Once localized, GPI-APs may be released by phospholipases or internalized by endocytosis (Ellis et al. 2010).

In Arabidopsis, homologs of all five subunits of the mammalian and yeast GPI-T complex—PIG-K/GPI8, GPAA1/GAA1, PIG-S/GPI17, PIG-T/GPI16, and PIG-U/GAB1—have been identified (Ellis et al. 2010). The catalytic subunits, AtGPI8 and AtPIG-S, have been studied in detail (Bundy et al. 2016; Liu et al. 2016; Desnoyer et al. 2020). Mutations in AtGPI8 cause severe developmental defects, including impaired cell wall integrity, growth, and fertility, due to disrupted GPI-AP biosynthesis and localization (Bundy et al. 2016). AtGPI8 is required for GPI-AP biosynthesis and proper localization of GPI-APs (Bundy et al. 2016). The *gpi8* mutant also shows defective pollen tube reception and altered membrane organization (Liu et al. 2016). AtPIG-S is critical for embryogenesis and pollen tube emergence and growth (Desnoyer et al. 2020). While loss-of-function mutations in these genes do not impair female gametophyte function, they disrupt LRE localization in the filiform apparatus (FA) of synergid cells, underscoring their role in GPI anchor addition (Liu et al. 2016; Desnoyer et al. 2020).

GPI-APs frequently localize to detergent-resistant membrane (DRM) microdomains, or lipid rafts, enriched in sterols and sphingolipids that facilitate protein sorting and signaling. Proteomic analyses of Arabidopsis DRM fractions have identified several GPI-APs—30, 59, and 163 in the respective studies by Borner et al. 2003; Elortza et al. 2003, 2006; Takahashi, Kawamura and Uemura, 2016, supporting their roles in key cellular processes within these microdomains.

The development of GPI-AP prediction tools has advanced our understanding of their prevalence in plant genomes. The first tool, “big-Pi predictor,” was based on metazoan and protozoan features (Eisenhaber, Bork and Eisenhaber, 1998, 1999), and later adapted for plants using validated and predicted Arabidopsis GPI-APs (Eisenhaber et al. 2003), identifying ∼245 GPI-APs (∼1% of the genome). A similar estimate (248) was made using GPT, a plant-specific tool based on proteomic analysis of PI-PLC–released GPI-APs from Arabidopsis callus tissue (Borner et al. 2003; Elortza et al. 2003).

Bioinformatic and proteomic approaches have identified pollen-expressed GPI-APs. Genome-wide microarray analysis in Arabidopsis revealed 47 pollen-expressed GPI-APs (Lalanne et al. 2004). Later, 248 predicted GPI-APs (Borner et al. 2003) and two PLD-identified ones (Elortza et al. 2006), were examined in laser-capture microdissected gametophyte cell datasets (Wuest et al. 2010). Of the 207 GPI-APs present on the array, 142 were expressed in at least one gametophyte cell type, with 23 male-specific and 48 female-specific (Desnoyer and Palanivelu, 2020).

Despite progress, most plant GPI-APs remain biochemically uncharacterized. A key exception is Arabidopsis SKU5, involved in directional root growth and confirmed as GPI-anchored via PI-PLC sensitivity and subcellular localization (Sedbrook et al. 2002). This lack of characterization is more acute in reproductive tissues, where relevant cells are few, small, and deeply embedded. While pollen is abundant for protein studies, isolating enough female gametophytic tissue is difficult. For example, LRE, essential for female gametophyte function, is expressed only in the two synergid cells per ovule.

Genetic evidence supports classifying LRE as a GPI-AP. Using the big-Pi plant predictor, we identified all typical GPI anchor domains in Arabidopsis LRE, including the GAS and ω sites (Eisenhaber et al. 2003). An LRE-cYFP fusion localized to the FA, a membrane-rich structure key for pollen tube reception. Deleting the GAS domain disrupted this localization and reduced FA signal, indicating impaired GPI anchoring. Similarly, removing predicted ω sites (Serine-139 and Alanine-141) reduced, but did not eliminate, FA localization (Liu et al. 2016). Mutations in transamidase components AtGPI8 and AtPIG-S also diminished LRE-cYFP localization (Liu et al. 2016; Desnoyer et al. 2020).

Unexpectedly, while deletion of the GAS domain or both predicted ω sites (one canonical and one cryptic) reduced the localization of LRE-cYFP at the FA, these changes did not impair its function in pollen tube reception and fertility (Liu et al. 2016). These results highlighted the complexity of GPI anchoring and revealed that low levels of LRE, even if incompletely GPI-anchored, may be sufficient to support its function in reproduction. Still, direct biochemical evidence in support of LRE’s GPI anchoring had been lacking. In this study, we address this using enzymatic assays on wild-type and variant LRE-cYFP proteins, providing direct evidence of GPI anchoring. Our results confirm the conservation and functional importance of GPI anchor domains in plant proteins.

## Results

### Plant-Specific Prediction Tools Accurately Identify Canonical and Cryptic ω Sites in LORELEI, a Putative GPI-Anchored Protein Expressed in the Synergid Cells of the Female Gametophyte

Previous analysis showed that deleting the top two predicted ω sites (Serine-139 and Alanine-141) significantly reduced—but did not eliminate—LRE-cYFP localization to the FA, suggesting existence of additional cryptic ω sites in LRE (Liu et al. 2016) (Fig. 1). Therefore, we performed further *in silico* analysis using metazoan-specific and big-Pi plant predictors to identify additional ω sites that could potentially contribute to GPI anchoring.

Analysis of the LRE sequence using the metazoan-specific predictor (Eisenhaber, Bork and Eisenhaber, 1998), revealed a different set of predicted ω-site residues than those identified by the plant-specific predictor (Fig. 1A, 1B). Notably, the plant predictor’s top-ranked ω site was ranked third by the metazoan tool and was the only overlap among their top three sites. To test the metazoan predictions, we deleted its top three ω sites in the *pLRE::LRE-cYFP* construct (designated *pLRE:LRE-cYFPΔ3ω*; Supplementary Fig. S1, Fig. 1A, 1B) and introduced it into the *lre-7/lre-7* mutant (Liu et al. 2016).

In synergid cell FA of these transformants, we assessed YFP signal as a proxy for PM localization of the LRE-cYFP fusion protein (Fig. 1C, D), following established methods (Liu et al. 2016). The LRE-cYFPΔ3ω variant showed no obvious mislocalization and only a modest reduction in the FA YFP signal (∼52%, Fig. 1H) compared to wild-type LRE-cYFP (∼68%, Fig. 1E) or LRE-cYFPΔ1ω (∼62%, Fig. 1F). Since both 1ω and 3ω deletions yielded similar FA signal levels, and serine-139 is the only shared missing residue, alanine-138 and aspartic acid-142 are unlikely ω-sites. Consistent with results reported in Liu et al. 2016, LRE-cYFPΔ2ω showed a marked reduction in FA signal (∼28%, Fig. 1G). Similarly, deleting all five top-predicted ω-sites using plant and metazoan predictors (*pLRE:LRE-cYFPΔ5ω*; Supplementary Fig. S1, Fig. 1B) caused a comparable defect (∼30% FA signal; Fig. 1I). These results suggest serine-139 and alanine-141 are the top ω-site candidates, with alanine-141 likely the primary site for GPI anchoring, as constructs lacking it showed consistent FA signal reduction, whereas deletion of serine-139 alone had little effect.

Since the newly generated *pLRE:LRE-cYFPΔ3ω* and *pLRE:LRE-cYFPΔ5ω* constructs showed localization patterns similar to *pLRE:LRE-cYFPΔ3ω*, we tested whether they could also rescue the pollen tube reception and seed set defects in *lre-7/lre-7* mutants, as previously shown for LRE-cYFPΔ2ω (Liu et al. 2016). Using a GUS staining assay (Tsukamoto et al. 2010), we found that both constructs fully complemented the pollen tube reception defect in *lre-7* female gametophytes (Supplementary Fig. S2A). Seed set was also restored to wild-type levels in *lre-7/lre-7* plants expressing either construct (Supplementary Fig. S2B), indicating that loss of 3ω or 5ω sites is as well tolerated as loss of 2ω sites.

Furthermore, when heterozygous transgenic plants carrying either *pLRE:LRE-cYFPΔ3ω* or *pLRE:LRE-cYFPΔ5ω* served as the female parent in crosses with wild type, progeny from these crosses showed increased transmission of *pLRE:LRE-cYFPΔ3ω* or *pLRE:LRE-cYFPΔ5ω* transgene, respectively (Supplementary Table S1). These results suggest that either transgene functionally complements LRE in the *lre-7* female gametophyte by rescuing defects in pollen tube reception and seed set (Supplementary Table S1). No such increase in transmission was observed when either of the transgenic plant was used as the male parent (Supplementary Table S1).

Taken together, these results demonstrate that deletion of many plausible ω-sites failed to yield additional insights beyond those already gained from the *pLRE:LRE-cYFPΔ2ω* construct. These findings also suggest that the big plant-Pi predictor outperforms the metazoan predictor in identifying the most functionally relevant ω-site in LRE. Moreover, we also concluded that further mutational analysis of ω-sites is unlikely to reveal new aspects of LRE’s function as a GPI-AP.

### LORELEI fused to cYFP Associates with the Detergent-Resistant Membrane Following Heterologous Expression in vegetative tissues

To directly test whether LRE is a GPI-AP, we used biochemical methods to assess its localization and sensitivity to GPI-AP-specific enzymatic treatments. Since LRE is mainly expressed in synergid cells within pistils (Liu et al. 2016) (Fig. 1C and 1D), we used pistils from *pLRE::LRE-cYFP* plants (Supplementary Fig. S1) to isolate *in planta*-expressed LRE protein. Total proteins (TP) from pistils were probed with anti-GFP, detecting LRE-cYFP bands in transgenic lines but not in non-transgenic *Col-0* plants (Fig. 2, panel 1), confirming LRE-cYFP expression in pistils. As a positive control, SKU5—a known GPI-AP (Sedbrook et al. 2002)—was similarly detected by anti-SKU5 (Fig. 2, panel 2).

**Fig. 2.**
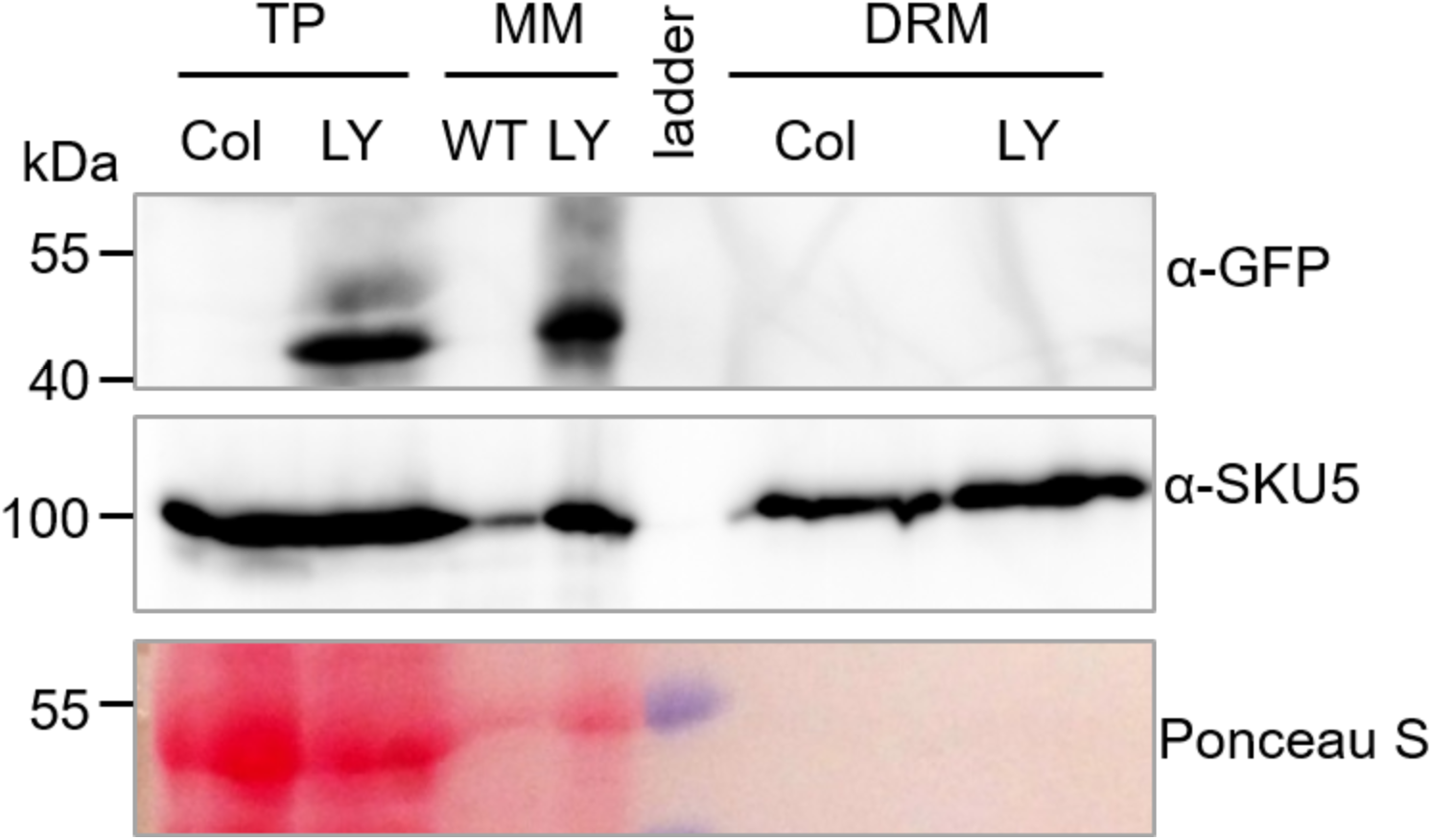
LRE-cYFP (LY) is present in total protein (TP) and microsomal membrane (MM) fractions but is undetectable in detergent-resistant membranes (DRM) from pistils containing the *pLRE::LRE-cYFP* transgene. Top panel: Western blot probed with anti-GFP (α-GFP) to detect LRE-cYFP. Middle panel: Western blot probed with anti-SKU5 (α-SKU5) to detect positive control SKU5, a known GPI-AP. Notably, Ponceau S staining, a widely used reversible stain to assess protein loading (Sander et al. 2019), likely showed a decreased signal in the MM fraction and no signal in the DRM fraction because these fractions are enriched for membrane components and may exclude most soluble proteins that Ponceau S typically detects. Col, pistils from non-transgenic wild-type Arabidopsis *Col-0* plants; LY, pistils from homozygous transgenic plants carrying the *pRBSC1A::LRE-cYFP* transgene; TP, total protein; MM, microsomal membrane protein; DRM, detergent-resistant membrane protein.

We isolated microsomal membrane proteins (MM) from total protein (TP) extracts and then extracted Triton X-114 detergent-resistant membranes (DRM) from the MM fraction for digestion with phosphatidylinositol-specific phospholipase C (PI-PLC), which cleaves the lipid moiety of GPI anchors (Doering, Englund and Hart, 1993) (Supplementary Fig. S3). In LRE-cYFP pistils, we detected SKU5 in both MM and DRM fractions (Fig. 2). LRE-cYFP was detected in the MM fraction, confirming membrane localization, but not in the DRM fraction (Fig. 2). This absence may reflect the restricted expression of LRE in synergid and egg cells of unpollinated pistils (Fig. 1C, D) (Liu et al. 2016; Wang et al. 2017).

To overcome low protein yield and obtain sufficient DRM for PI-PLC assays, we ectopically expressed the LRE-cYFP fusion in leaves using the *RBSC1A* promoter. The LRE-cYFP sequence used matched the one shown to rescue the *lre* mutant seed phenotype (Liu et al. 2016). After selecting homozygous lines with strong expression (*pRBSC1A::LRE-cYFP*; Supplementary Fig. S4), we used 14-day-old seedlings (excluding roots, as RBSC1A is mesophyll-specific) to isolate TP, MM, and DRM (Supplementary Fig. S3). SKU5 was detected in all fractions except in the *sku5* null mutant (Fig. 3). Notably, LRE-cYFP was detected in TP, MM, and DRM, confirming its presence in membrane fractions, including the detergent-resistant membrane from leaves (Fig. 3).

**Fig. 3.**
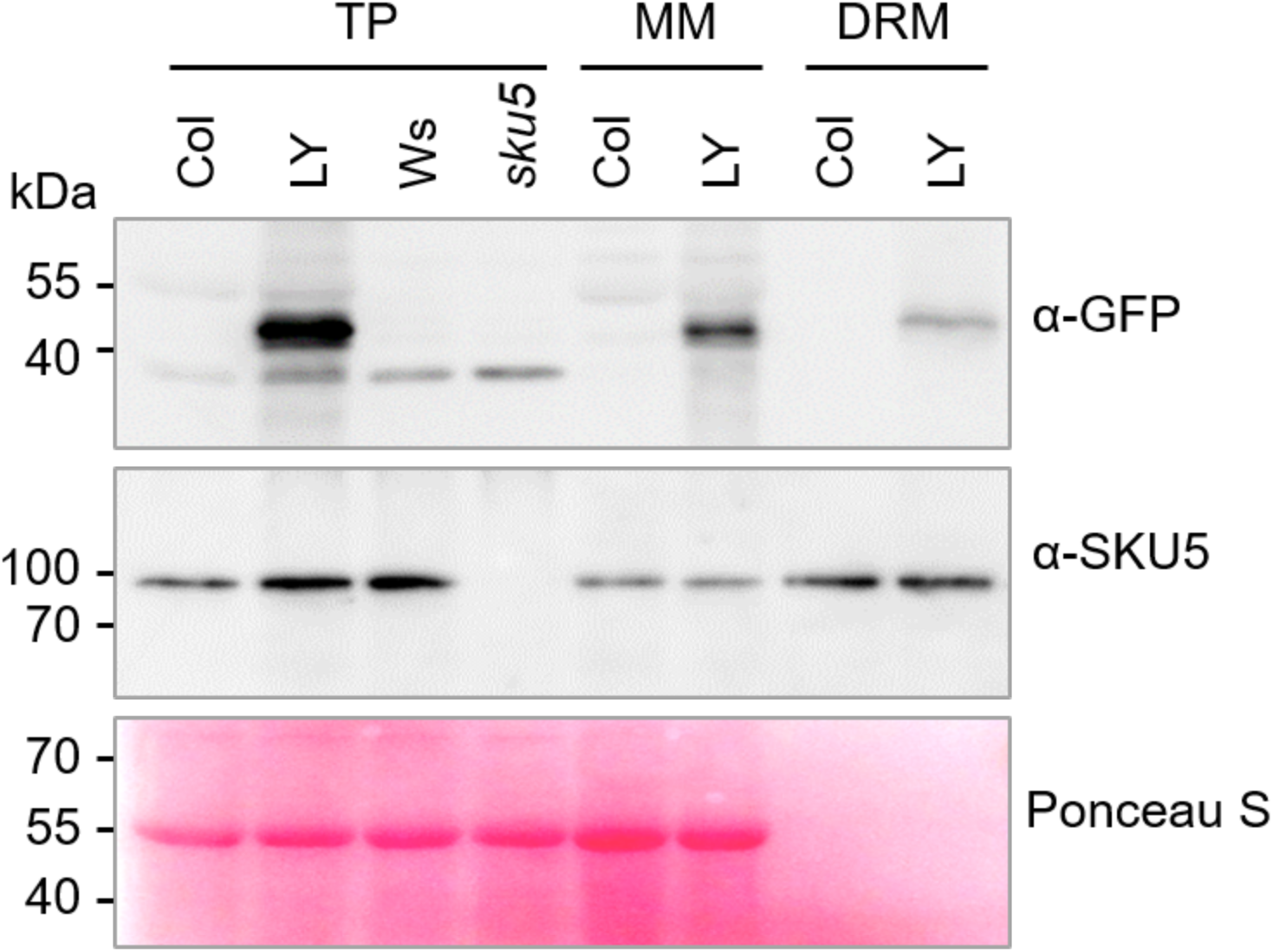
LRE-cYFP (LY) is present in TP, MM, and DRM fractions from seedlings expressing *pRBSC1A::LRE-cYFP*. Top panel: Western blot probed with anti-GFP (α-GFP) to detect LRE-cYFP in different fractions. Middle panel: Western blot probed with anti-SKU5 (α-SKU5) to detect positive control SKU5, a known GPI-AP. Bottom panel: Ponceau S staining to assess protein loading. Col/Ws, 14-days-old seedlings from non-transgenic wild-type Arabidopsis *Col-0* or *Ws* plants, which serve as background controls for the LY transgene and the *sku5* mutation, respectively; LY, 14-day-old seedlings from homozygous transgenic plants carrying the *pRBSC1A::LRE-cYFP* transgene; TP, total protein; MM, microsomal membrane protein; DRM, detergent-resistant membrane protein.

### LRE is a GPI-Anchored Membrane Protein

To test if LRE is a GPI-AP, we treated DRM fractions from *pRBSC1A::LRE-cYFP* seedlings with PI-PLC, which releases GPI-APs from DRMs into the aqueous phase (Doering, Englund and Hart, 1993) (Supplementary Fig. S3). We collected the upper (UP4, aqueous) and lower (LP4, DRM) phases from PI-PLC- and mock-treated samples (Supplementary Fig. S3). Immunoblotting with anti-GFP detected LRE-cYFP in these fractions. To monitor protein recovery efficiency from the upper phase after the PI-PLC treatment, we added a fixed amount of BSA, a soluble protein, as an internal control. Ponceau S staining of UP4 fractions confirmed a comparable BSA recovery across all aqueous phase samples (UP4, Fig. 4), indicating comparable recovery of SKU5 or LRE-cYFP proteins from non-transgenic *Col-0* wild type and transgenic samples. As expected, SKU5, a known GPI-AP, accumulated predominantly in the aqueous phase only after PI-PLC treatment, remaining in the DRM phase otherwise (Fig. 4). Similarly, LRE was detected in the aqueous phase exclusively after PI-PLC treatment, confirming that LRE is a GPI-AP (Fig. 4).

**Fig. 4.**
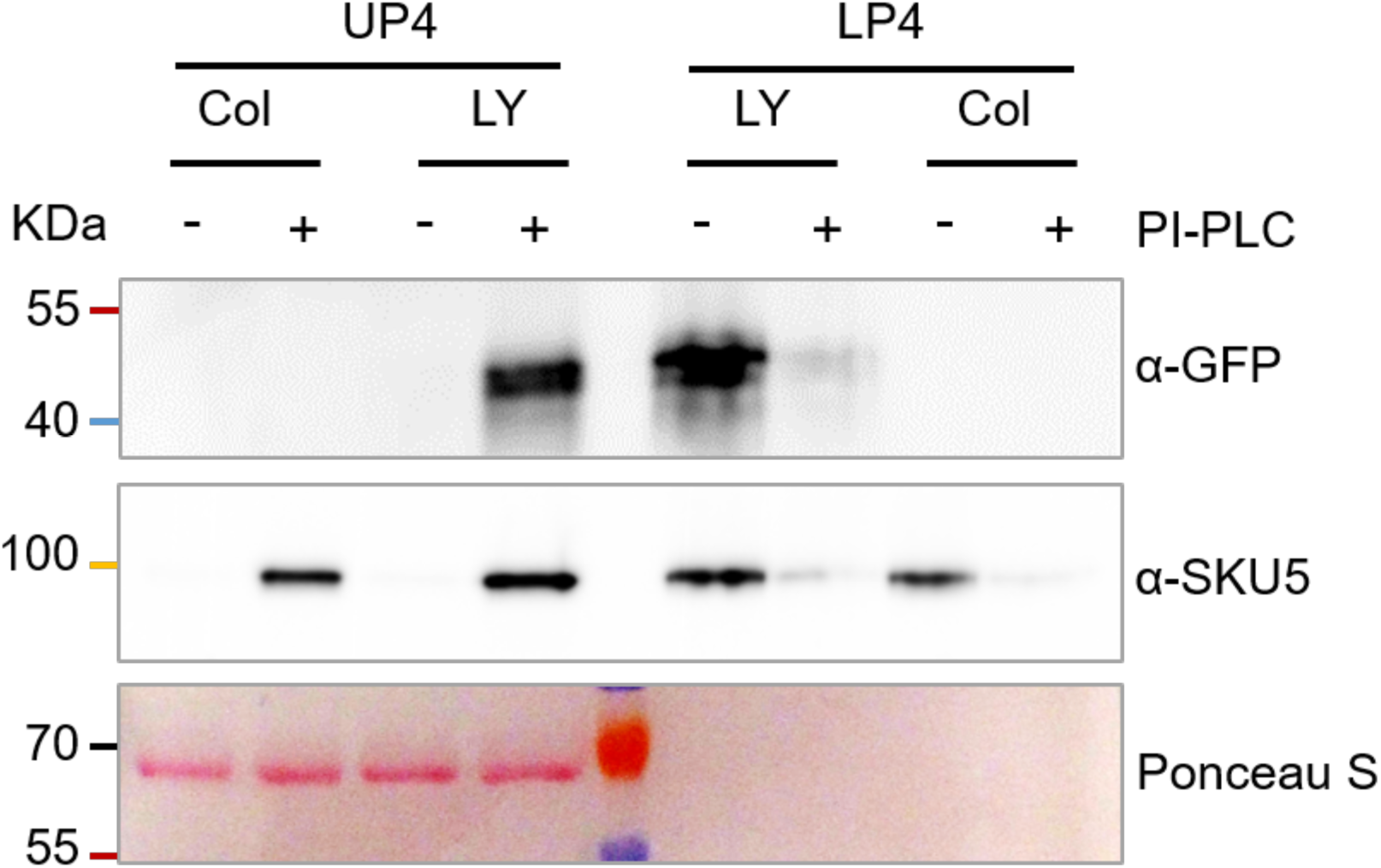
LRE-cYFP (LY) is released from the DRM fraction into the aqueous phase after the PI-PLC treatment. Top panel: Western blot probed with anti-GFP (α-GFP) to detect LRE-cYFP. Middle panel: Western blot probed with anti-SKU5 (α-SKU5) to detect SKU5, a known GPI-AP, as a positive control. Bottom panel: Ponceau S staining is used to assess protein recovery after precipitation from the aqueous phase, as indicated by externally added BSA protein. UP4, the upper phase after the fourth PI-PLC treatment (Supplementary Fig. S3), represents the aqueous phase following phase separation. LP4, the lower phase after the fourth PI-PLC treatment, represents the membrane phase following phase separation. Col, 14-day-old seedlings from non-transgenic wild-type Arabidopsis *Col-0* plants. LY, 14-day-old seedlings from homozygous transgenic plants carrying the *pRBSC1A::LRE-cYFP* transgene.

### Functional Analysis of GPI-Anchoring Domains in LRE Localization and Membrane Association

In mammalian cells, a GPI transaminase recognizes the ω of a proprotein, cleaves the GAS domain, attaches a GPI moiety to the ω site, forms a nascent GPI-AP in the endoplasmic reticulum (ER). This nascent protein matures during transport to the PM (Kinoshita, 2020). A similar process has been proposed for plant GPI-AP biosynthesis (Ellis et al. 2010).

Our previous genetic analysis showed that deleting two ω sites (predicted and a cryptic site) or the GAS domain (ΔGAS) at the C-terminus of the LRE protein caused a dramatic change in the localization of reporter fusion proteins in the FA of synergid cells (Liu et al. 2016) (Fig. 1), consistent with the prediction that LRE is a GPI-AP. To investigate the importance of these domains in GPI-anchoring of LRE as determined by the PI-PLC-based assay described above, we generated variant constructs lacking either ω sites or GAS domains or replacing both with a transmembrane domain (TM) of FERONIA receptor-like kinase under the *35S promoter* for ectopic expression in leaves (*p35S::LRE-cYFPΔ2ω, p35S::LRE-cYFPΔGAS*, and *p35S::LRE-cYFP-TM,* Supplementary Fig. S4). We selected LRE-cYFPΔ2ω to represent the ω-site deletion constructs, as it resulted in the most pronounced reduction in LRE localization to the FA of synergid cells (Fig. 1J), and LRE-cYFPΔ5ω did not differ significantly from LRE-cYFPΔ2ω in this regard. Based on our localization alteration results described in Fig. 1 and in Liu et al. 2016, we predicted the following outcomes *in planta*: The LRE-cYFPΔ2ω variant lacks ω sites required for recognition by a transamidase, preventing the addition of a GPI anchor. As a result, it will fail to associate with the PM and remain insensitive to PI-PLC treatment. Conversely, the LRE-cYFPΔGAS variant would lack the GAS domain and GPI-anchor, rendering it non-membrane bound and released from the cell, thereby unavailable as a substrate for PI-PLC treatment. In contrast, the LRE-cYFP-TM variant would remain membrane-bound and be insensitive to PI-PLC treatment due to the presence of a transmembrane domain.

These constructs were transiently expressed in *Nicotiana benthamiana* leaf epidermal cells to validate *in planta* expression and assess subcellular localization. As observed in the FA of synergid cells (Liu et al. 2016), LRE-cYFP localized predominantly to cell boundaries, likely the PM. This was supported by its co-localization with the PM marker PIP2A-mCHERRY (Fig. 5A, bottom panel). To confirm this, LRE-cYFP and PIP2A-mCHERRY were co-expressed in tobacco leaves. The merged YFP and mCHERRY signals at the cell boundaries confirmed PM localization (Fig. 5B). LRE-cYFP-TM showed a similar pattern and co-localization with PIP2A-mCHERRY (Fig. 5A–B).

**Fig. 5.**
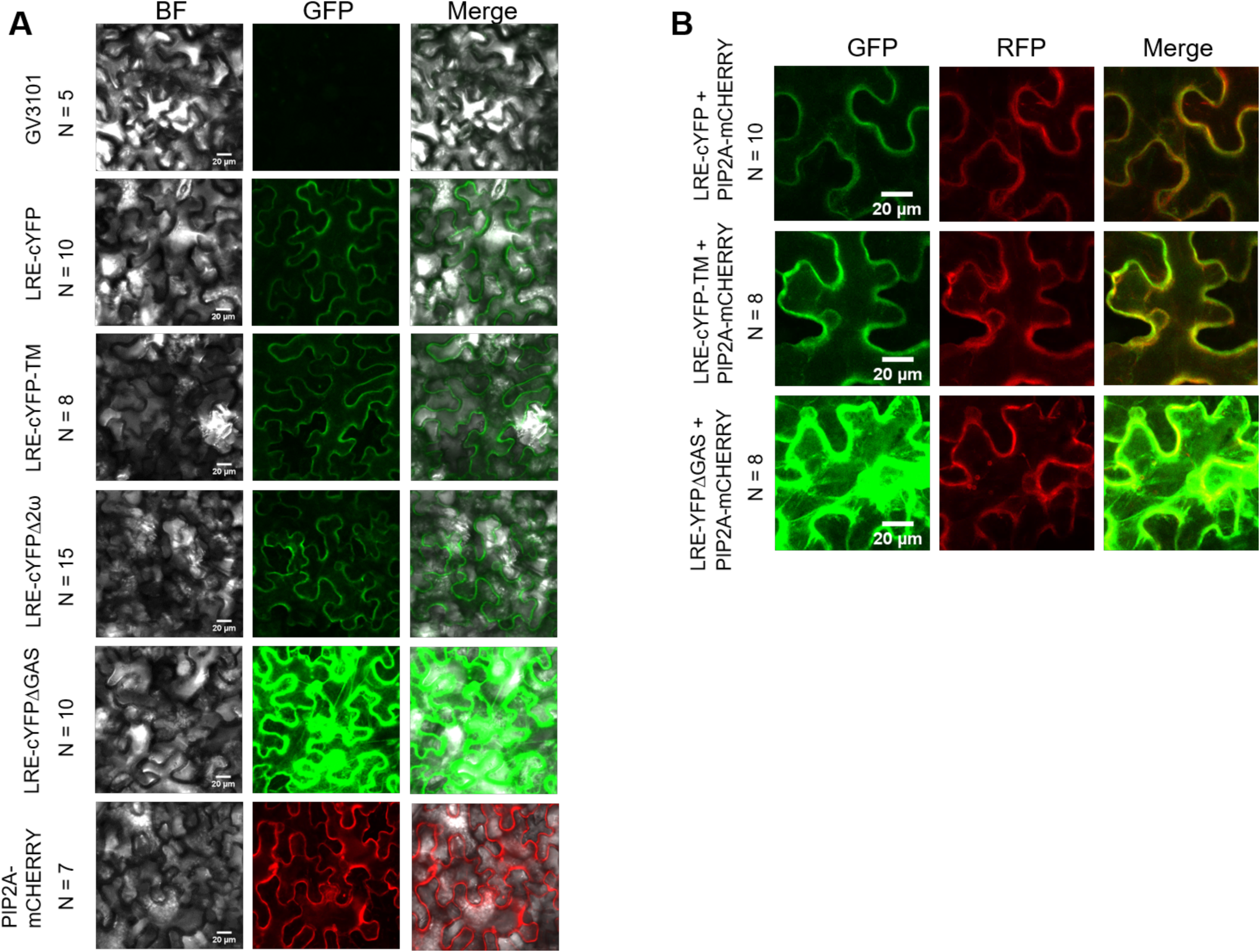
Transient expression of *AtLRE* variants under the control of the *35S* promoter in *Nicotiana benthamiana* leaves. (A) Transient expression of *AtLRE-cYFP*, *AtLRE-cYFP-TM*, *AtLRE-cYFPΔ2ω*, *AtLRE-cYFPΔGAS*, or *PIP2A-mCHERRY*, a plasma membrane marker, in *N. benthamiana* leaves. Each image panel is a z-stack projection of nine optical sections from infiltrated leaves. (B) Co-expression of selected *AtLRE* variants (*AtLRE-cYFP*, *AtLRE-cYFP-TM*, or *AtLRE-cYFPΔGAS*) with *PIP2A-mCHERRY*, a plasma membrane marker, in *N. benthamiana* leaves. Each image panel is a z-stack projection of nine optical sections from infiltrated leaves. BF, bright field; GFP, green fluorescence protein imaging channel; RFP, red fluorescence protein imaging channel. Scale bar, 20 µm.

LRE-cYFPΔ2ω and LRE-cYFPΔGAS localized to the PM or PM-like structures (Fig. 5A). However, LRE-cYFPΔGAS showed broader subcellular distribution, extending into the cell interior or extracellular space beyond the membrane (Fig. 5A). In co-localization with PIP2A-mCHERRY, much of the LRE-cYFPΔGAS signal appeared around the PM, unlike the tight membrane association of LRE-cYFP-TM (Fig. 5B), indicating that a notable portion of LRE-cYFPΔGAS is not retained in the PM.

### The ω Sites and GAS Domain are Critical for GPI Anchor Attachment to LORELEI

After characterizing expression and localization of LRE variants, we assessed the role of specific domains in GPI-anchoring using the PI-PLC assay. Constructs were transformed into wild-type *Col-0* plants to generate stable transgenic lines. In leaf tissues, all constructs were detectably expressed with localization patterns largely similar to those seen in *Nicotiana benthamiana*. LRE-cYFP and LRE-cYFP-TM localized mainly to the cell periphery, consistent with PM association (Supplementary Fig. S5). LRE-cYFPΔ2ω and LRE-cYFPΔGAS also showed peripheral localization but with broader distribution into the cell interior or extracellular space, suggesting reduced membrane association.

Seedlings (above-ground tissues from 2-week-old plants) were used to isolate TP, MM, and DRM fractions for the PI-PLC assay. Since we are comparing different constructs, it is essential to load comparable levels of transgenic protein, as individual transgenic lines may exhibit different expression levels due to position effects from the transgene insertion site in the genome or copy number variations. To control this, total proteins were isolated, and cYFP-tagged protein levels were assessed by western blot using anti-GFP antibodies. This enabled normalization of LRE input for DRM fraction isolation (Supplementary Fig. S6A).

After three rounds of Triton X-114 treatment, the purified DRM (Supplementary Fig. S3) was treated with either PI-PLC or a mock buffer (without PI-PLC). Probing the fractions with SKU5 confirmed the successful isolation of distinct fractions and that the PI-PLC treatment was effective (Fig. 6). A similar pattern was observed with LRE-cYFP. Before PI-PLC treatment, the LRE-cYFP remained in the DRM (Fig. 6). After PI-PLC treatment, the LRE-cYFP was released to the aqueous phase (Fig. 6), consistent with the behavior reported above (Fig. 2). As expected, LRE-cYFP-TM was insensitive to PI-PLC treatment and remained in the DRM (Fig. 6).

**Fig. 6.**
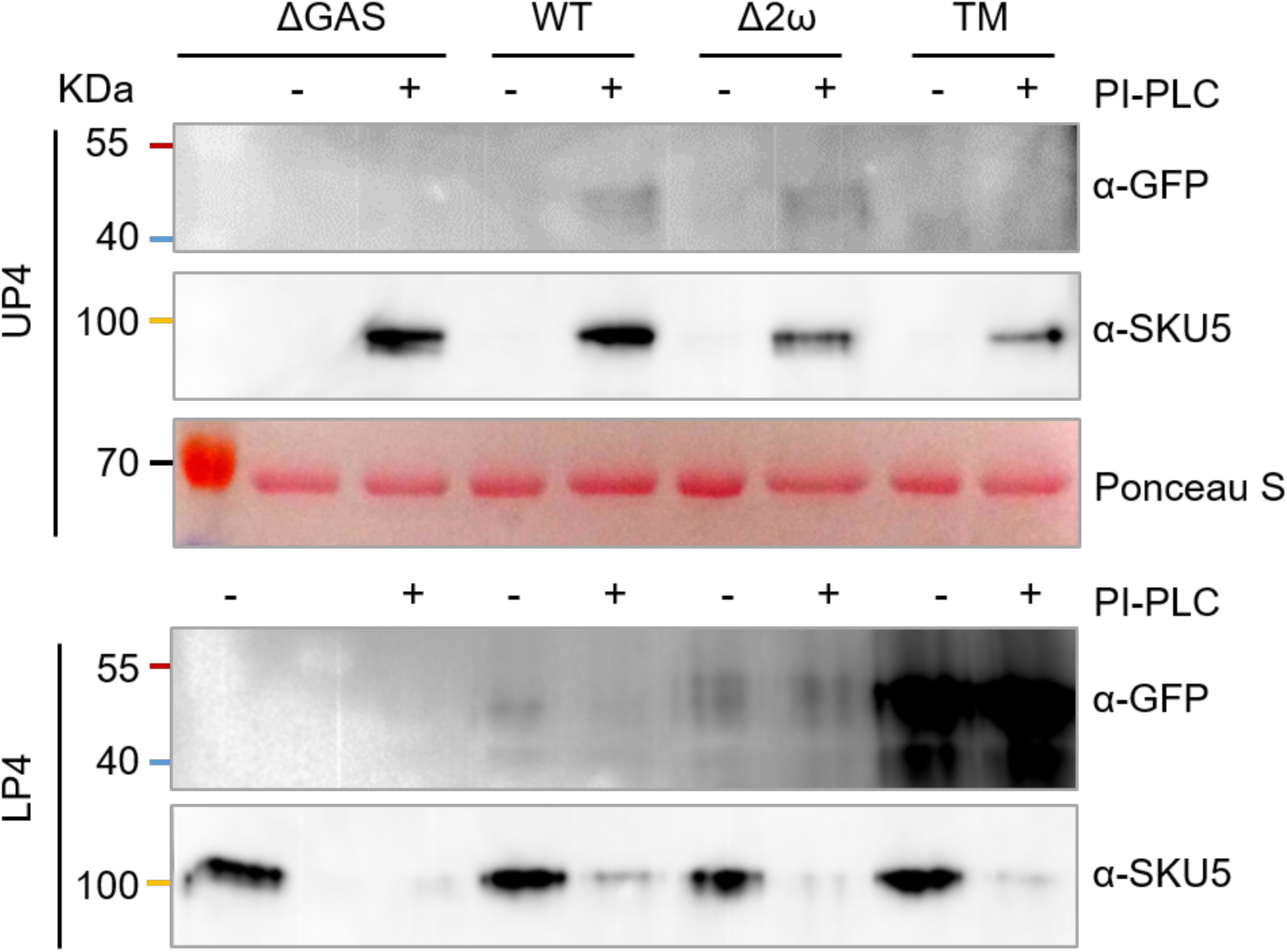
The GAS domain and omega sites in the C-terminus of LRE are involved in GPI anchoring LRE to the membrane. The top three panels show western blot analysis using protein fractions from the UP4 phase (Supplementary Fig. S3). The first panel shows a western blot probed with a GFP antibody (α-GFP) to detect LRE-cYFP. The second panel shows a western blot probed with a SKU5 antibody (α-SKU5) to detect the positive control SKU5, a known GPI-AP, and the third panel shows Ponceau S staining to assess protein recovery after precipitation from the aqueous phase indicated by externally added BSA protein. The bottom two panels show western blot analysis using protein fractions from the LP4 phase (Supplementary Fig. S3). The first panel shows a western blot probed with a GFP antibody (α-GFP) to detect LRE-cYFP, and the second panel shows a western blot probed with a SKU5 antibody (α-SKU5) to detect the positive control SKU5, a known GPI-AP. UP4: Upper phase after the 4th round of treatment, representing the aqueous phase of the two separated phases following PI-PLC treatment. LP4: Lower phase after the 4th round of treatment, representing the membrane phase of the two separated phases following PI-PLC treatment. ΔGAS: Protein from seedlings carrying a *p35s::LRE-cYFP ΔGAS* transgene. WT: Protein from seedlings carrying a *p35s::LRE-cYFP-WT* transgene. Δ2ω: Protein from seedlings carrying a *p35s::LRE-cYFP Δ2ω* transgene.

In the case of LRE-cYFPΔ2ω, we predicted that it would not receive a GPI-anchor, remain in the DRM, and be insensitive to PI-PLC treatment. While a portion of LRE-cYFPΔ2ω was present in the DRM after PI-PLC treatment, indicating that the deletion of the two predicted ω sites affected GPI anchoring, some LRE-cYFPΔ2ω was also detected in the aqueous phase (Fig. 6). This suggests that additional cryptic ω sites in the LRE protein sequence, or an as-yet undefined mechanism, may contribute to the continued membrane association of a portion of these variant proteins. It’s also possible that unknown domains with redundant functions with ω sites are present, albeit less efficiently than the predicted sites. Nonetheless, our biochemical analysis confirmed that the predicted ω sites are critical for GPI anchoring in LRE.

Regarding LRE-cYFPΔGAS, it was absent from both the upper and lower phases, regardless of the PI-PLC treatment (Fig. 6). This absence of LRE-cYFPΔGAS in the DRM was consistent with our prediction that deleting the GAS domain prevents LRE from receiving a GPI anchor. However, the absence of LRE-cYFPΔGAS in the aqueous fraction after PI-PLC treatment raised whether it might have been part of earlier supernatant fractions during the DRM isolation procedure due to its non-membrane-associated nature. To investigate this possibility, we examined the aqueous fractions (SUP, UP1, UP2, and UP3) collected during the DRM isolation procedure (Supplementary Fig. S3). We found that LRE-cYFPΔGAS was present in the SUP and UP1 fractions but was absent in subsequent steps (Supplementary Fig. S6B). As a control for DRM isolation efficiency, we used SKU5 as a positive GPI-AP control and UGPase as a marker for soluble fractions. These findings demonstrate that LRE-cYFPΔGAS is not part of the (DRM) and underscore the critical role of the GAS domain in facilitating proper GPI anchoring of LRE in plants.

As an alternative approach to examining the *cis* domains (ω site and GAS domain) important for GPI anchoring, we also evaluated the role of transacting factors in this process. Previous research demonstrated that loss-of-function mutants in *GPI8*, a key component of the transamidase complex responsible for GPI anchoring on GPI-APs, disrupt LRE localization in the FA of synergid cells, presumably due to the loss of GPI anchoring (Bundy et al. 2016; Liu et al. 2016). To investigate whether the transamidase complex affects LRE protein production *in planta*, we introduced LRE-cYFP into the *gpi8-1*, the weaker allele tested in Bundy *et al* 2016, as the *gpi8-2* allele tested in Liu et al. 2016 is lethal. As a negative control, we separately introduced LRE-cYFP, which is expected to be not a substrate of the transamidase complex due to the presence of a transmembrane domain, into the *gpi8-1* background. As predicted, no size shift was observed in the LRE-cYFP-TM bands between *Col-0* wildtype and *gpi8-1* backgrounds (Fig. 7), consistent with LRE-cYFP-TM being insensitive to GPI-8. In contrast, LRE-cYFP exhibited prominent higher molecular weight bands in the *gpi8-1* background compared to *Col-0* wild-type plants (Fig. 7). Although not experimentally validated, based on the size of the larger band(s), (Supplementary Table 2),we surmise that it likely corresponds to the unprocessed LRE proprotein with the GAS domain still present. Collectively, these findings involving using the *cis* domains and *trans*-acting factors critical for GPI anchoring in GPI-APs support the conclusion that the ω sites and the GAS domain within the LRE protein sequence are crucial for proper GPI-anchor attachment to LORELEI.

**Fig. 7.**
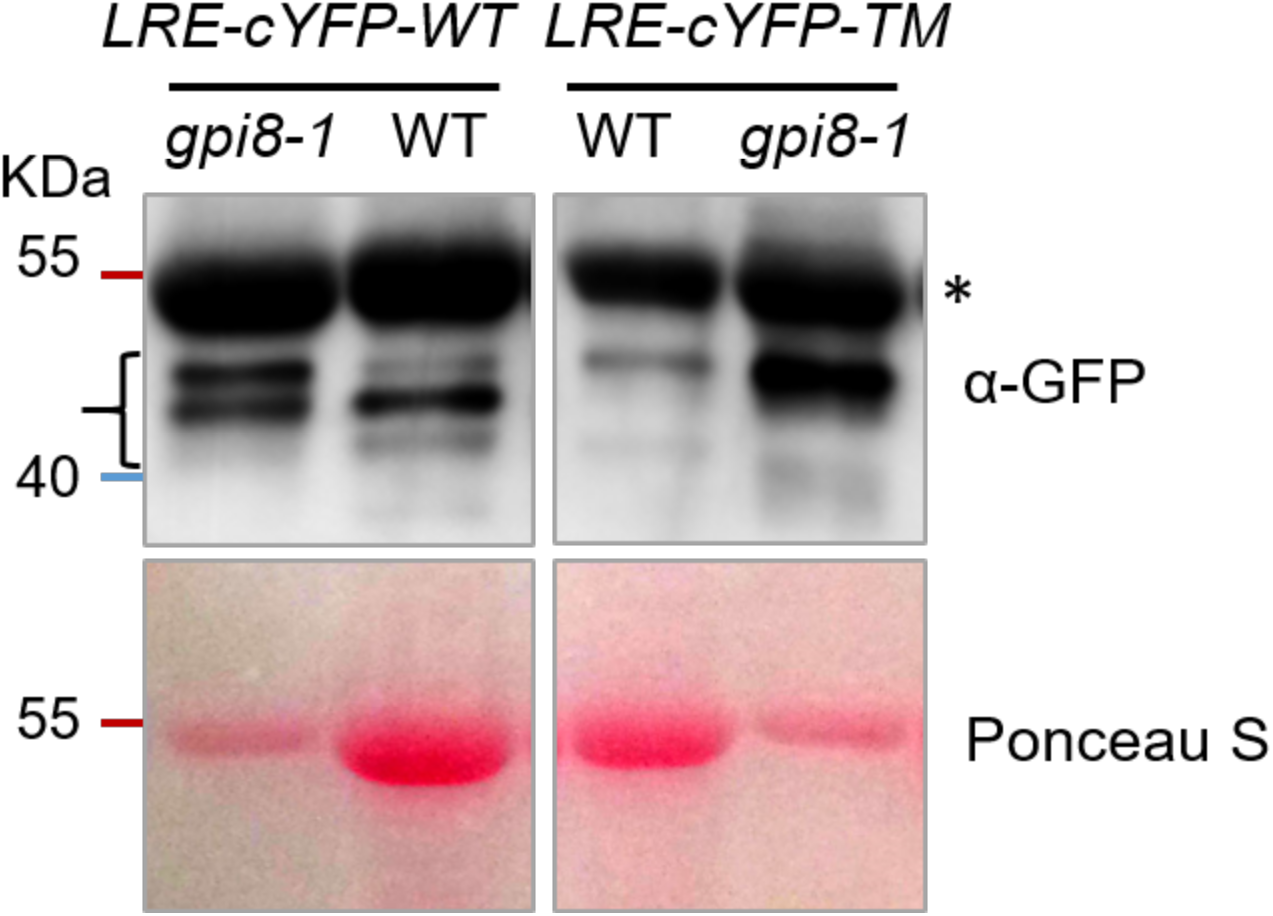
The loss-of-function allele of GPI8 (*gpi8-1*), a key component of the Arabidopsis transamidase complex, results in larger LRE proteins. The top left panel shows a western blot probed with GFP antibody (α-GFP) using total protein extracted from wild-type *Col-0* (WT) or *gpi8-1* mutant plants carrying a *p35S::LRE-cYFP-WT* transgene. The top right panel shows a western blot probed with GFP antibody (α-GFP) using total protein from WT or *gpi8-1* mutant plants carrying a *p35S::LRE-cYFP-TM* transgene. The bottom panels show Ponceau S staining for protein loading. Based on size estimation, the LRE-cYFP bands are indicated by a bracket. Unknown non-specific bands, present in all plants assayed in this experiment, are marked with an asterisk (*).

## Discussion

In this study, we present the first biochemical evidence that LRE is a GPI-AP, resolving longstanding questions about its membrane association. Although LRE is expressed in pistils, it was undetectable in the DRM fraction, likely due to its expression in just four ovular cells (two synergids, one egg, one central). By ectopically expressing LRE in vegetative tissues, we obtained sufficient protein for biochemical analysis and confirmed it is a GPI-AP. In leaf cells, LRE localizes to the cell periphery and associates with the DRM fraction, is cleavable by PI-PLC, and requires specific *cis*-domains for proper GPI anchoring. These findings clarify LRE’s biochemical properties and lay the groundwork for future studies of GPI-anchored proteins in plant development.

### GPI-anchored Protein Prediction in Plants

We introduced point mutations at cryptic ω sites in the LRE sequence, predicted by two independent algorithms, to assess their effects on LRE localization and function. Neither LRE-cYFPΔ3ω (ω sites identified by the metazoan predictor) nor LRE-cYFPΔ5ω (ω sites identified by both predictors) showed greater disruption than LRE-cYFPΔ2ω (ω sites identified by the plant predictor). Based on the amino acids deleted in each construct, Alanine-141 is likely the primary ω site, as two constructs that resulted in YFP signal reduction in FA included loss of Alanine-141 (Fig. 1B, 1J). While the combined deletion of Serine-139 and Alanine-141 in the 2ω construct may contribute to this loss, the 3ω construct—which also lacks Serine-139—was fully functional, suggesting Alanine-141 plays the key role. Alanine-135, predicted as the fourth ω site by the metazoan predictor, had no effect on localization, highlighting the greater accuracy of the big-Pi plant predictor for plant sequences. Definitive identification of the ω site will require protease digestion of mature LRE followed by tandem mass spectrometry (Eisenhaber et al. 2003).

### LRE is a GPI-Anchored Protein

Although LRE was previously suggested to be membrane-associated (Liu et al. 2016), its classification as a GPI-AP had not been biochemically confirmed. Using *in planta* expression, subcellular fractionation, and PI-PLC treatment, we showed that LRE exhibits hallmark features of GPI-APs. Immunoblotting detected LRE in total protein (TP), microsomal membrane (MM), and DRM fractions, consistent with membrane association. Crucially, PI-PLC treatment solubilized LRE from DRM into the aqueous phase—similar to SKU5, a known GPI-AP—confirming its GPI-anchoring. While ∼250 GPI-APs have been predicted in Arabidopsis (Borner et al. 2003; Desnoyer and Palanivelu, 2020), LRE is now one of the few that have been biochemically confirmed as a GPI-AP.

To facilitate detection, LRE was fused to cYFP, allowing use of a monoclonal GFP antibody. The reporter did not alter localization or processing, and the fusion protein remained a transamidase substrate. Its pH insensitivity also enabled reliable comparison across variants. These results demonstrate cYFP’s suitability for GPI-AP biochemical analysis.

### Functional Importance of the ω Sites and GAS Domain in GPI Anchoring

Subcellular localization in *Nicotiana benthamiana* leaves revealed the critical roles of ω sites and the GAS domain in GPI anchoring. This system effectively validated LRE constructs and enabled visualization of LRE and its variants. Localization patterns in *Nicotiana benthamiana* largely matched those in Arabidopsis, supporting its utility. As expected, deleting ω sites partially retained membrane localization, while GAS domain deletion disrupted cell-peripheral localization. These patterns were confirmed in stable lines and biochemical assays. The Δ2ω variant retained some DRM association after PI-PLC treatment, suggesting partial anchoring via cryptic ω sites. In contrast, the ΔGAS variant lost membrane association and accumulated in soluble fractions, underscoring the GAS domain’s essential role—similar to GP63 in Leishmania, which lacks a functional GPI anchor (McGwire et al. 2002).

### Role of Transamidase Complex in LRE Processing

We investigated the role of trans-acting factors in LRE processing by expressing LRE-cYFP in the *gpi8-1* mutant background. This led to prominent higher molecular weight bands, likely representing unprocessed LRE, providing genetic evidence that GPI8 is essential for LRE maturation and GPI anchoring. A transmembrane-anchored LRE version (LRE-cYFP-TM) was unaffected by PI-PLC treatment or the *gpi8-1* mutation, confirming that the observed effects were specific to GPI anchoring. These findings align with prior characterizations of GPI8 in Arabidopsis (Bundy et al. 2016) and support the identification of LRE as a GPI8 substrate (Liu et al. 2016). Given its small size (41–45 kDa), LRE is particularl y useful for detecting size shifts caused by mutations in LRE or transamidase components. Such changes are less apparent in larger GPI-APs like SKU5 (∼90 kDa). LRE thus serves as an effective model for studying mutations affecting GPI anchor addition. It would be valuable to test whether similar processing defects occur in mutants of other transamidase subunits, such as PIG-S (Desnoyer et al. 2020).

### The Functional Implications of Loss of GPI Anchoring in LRE Variants

Previous studies showed that mutant LRE proteins lacking GPI anchor domains could still rescue pollen tube reception and seed set defects (Liu et al. 2016), raising the question of why these conserved domains appear dispensable for function. One explanation is that mutant proteins may accumulate at the FA at sufficient levels to trigger reception, despite impaired membrane anchoring. This assumes that the loss of these domains results in a partial reduction in membrane anchoring (as seen in the Δ2ω variant) or a complete release of the protein from the PM (as observed in the ΔGAS variant). However, direct biochemical evidence supporting this possibility has been lacking.

Our biochemical analysis supports this model. The Δ2ω variant showed reduced GPI anchoring, with protein detected in both soluble and membrane fractions after PI-PLC treatment. In contrast, the ΔGAS variant was mostly found in the soluble phase, indicating near-complete loss of membrane association. These results confirm that the ω sites and GAS domain are essential for GPI anchoring, yet also suggest that mislocalized LRE variants may remain functional if present—even at low levels—in extracellular spaces where they can engage their targets.

### Novel Contributions and Broader Implications

This study makes several key contributions. First, we provide the first biochemical confirmation that LRE is a GPI-AP, resolving prior ambiguities and adding it to the short list of experimentally verified plant GPI-APs. Second, we define the cis-acting elements essential for anchoring—specifically, the ω sites and GAS domain. Third, we establish a functional connection between GPI transamidase activity and LRE processing, highlighting the significance of post-translational regulation in reproductive signaling.

These findings have broader implications. GPI-APs mediate cell-cell communication, especially in reproduction. Of the 207 Arabidopsis GPI-APs with expression data, 23 and 48 are male and female gametophyte-specific, respectively, with 21 linked to reproductive phenotypes (Desnoyer and Palanivelu, 2020). Yet, their biochemical study is limited by the small size of gametophytic cells. Here, we demonstrate that ectopic expression of LRE in leaves enables its proper processing and biochemical analysis, offering a viable strategy for characterizing other gametophyte-expressed GPI-APs. This approach sets the stage for future analysis of LRE family members (LLG1-3; Tsukamoto et al. 2010) and other plant GPI-APs (Borner et al. 2003).

## Conclusions

We demonstrate that LRE is a bona fide GPI-anchored protein, define the molecular determinants required for its anchoring, and link its processing to the GPI transamidase complex. These findings advance our understanding of peripheral membrane proteins in plant reproduction and provide a foundation for exploring their roles in cell-cell signaling and environmental responses. Given the conserved nature of GPI anchoring, similar mechanisms may govern the localization and function of other reproductive proteins across plants.

## Supporting information

Supplemental Data combined

## Acknowledgments and Funding

We thank Dr. John Sedbrook, Illinois State University, for providing the SKU5 antibody, WS, and *sku5* mutant seeds and for discussions on DRM isolation protocols. We thank Dr. Elena Shpak for sharing *gpi8-1* mutant seeds (Bundy et al. 2016). We thank Drs. Zhongguo Xiong, Shanshan Shi, Jennifer Noble, and Nathaniel Ponvert for their technical help. We thank Dr. Zhongguo Xiong for sharing the secondary antibody (goat-anti Rabbit HRP). We thank Drs. Rebecca Mosher, Mark Beilstein, David Baltrus, Karen Schumaker, and Ramin Yadegari for providing access to Azure c300, Western Blotting equipment, reagents, and their valuable discussions. We thank Drs. Choong-Hwan Ryu for assistance with the transient expression of various constructs in *Nicotiana benthamiana.* We thank Calvin J. Perkins for help with preparing Supplementary Fig. S2. G.H. was supported by funding from the BIO5 Institute and donors of the Undergraduate Biology Research Program (UBRP) at the University of Arizona. This work was supported by funding from the National Science Foundation (IOS 1146090) to R.P.

## Author Contributions

YW and RP designed the experiments. YW conducted protein fractionation, enzymatic assays, and Western blots. XL generated all constructs used in this study, performed complementation experiments and confocal imaging of ovules. ND analyzed the confocal images of ovules. ND and GH performed transient expression experiments in *Nicotiana benthamiana* and imaged subcellular localization of various LRE-cYFP variants in transgenic Arabidopsis leaves. YW and RP wrote the manuscript. All authors reviewed and approved the manuscript submitted for publication.

## Methods

### Plant material and growth conditions

The procedures for seed planting, seedling transplanting, and plant growth conditions followed the methods as described (Liu et al. 2016). All wild-type and transgenic plants used in this study were of the Columbia (*Col-0*) ecotype, except for *sku5*, which is in the Wassilewskija (*Ws*) background.

### Cloning transgenic constructs

To generate the Δ3ω and Δ5ω mutations in LRE-YFP under the native LRE promoter, site-specific changes were introduced into the LRE coding sequence via PCR using the *pLRE:LRE-cYFP* construct (Liu et al. 2016) as a template. Overlap PCR was used to assemble the full constructs, which were inserted into SpeI/AscI-linearized pLRE:LRE-cYFP plasmid using the In-Fusion HD Cloning Plus kit (Clontech; 638909), replacing the wild-type LRE-cYFP with the desired ω-site deletions.

To generate *pRBSCA1:LRE-cYFP*, the Arabidopsis *RBSCA1* promoter (1980 bp upstream of the start codon) was amplified from *Col-0* genomic DNA using primers 1759 (AGGCGGCCGCACTAGTTACCTTACGAGGAGCTTGAGCTTC, which included a *SpeI* restriction site) and 1760 (GCTCCATTGTTCTTCTTTACTCTTTGTGTGACTGAG). The *LRE-cYFP* sequence was amplified from *pLRE:LRE-cYFP* (Liu et al. 2016) plasmid using primers 1761 (GAAGAACAATGGAGCTGATATTATTATTCTTCTTTC) and 1702 (AGCTGGGTCGGCGCGCCGGAGGTCAAGTATTCTTTACACTTGGACACT, which included a *AscI* restriction site). The *pLRE:LRE-cYFP* plasmid was digested with SpeI and AscI to remove the original promoter, and the two PCR fragments were assembled into the gel-purified backbone using the In-Fusion HD Cloning Plus kit (Clontech; 638909), yielding the *pRBSCA1:LRE-cYFP* construct.

For *p35S:LRE-cYFP*, *p35S:LRE-cYFPΔ2ω*, *p35S:LRE-cYFPΔGAS*, and *p35S:LRE-cYFP-TM*, promoterless *LRE-cYFP*, *LRE-cYFPΔ2ω*, *LRE-cYFPΔGAS*, and *p35S:LRE-cYFP-TM* fragments, along with the T35 terminator, were first amplified from corresponding plasmids described in (Liu et al. 2016) using primers 1976 (CGGAGCTAGCTCTAGAAATGGAGCTGATATTATTATTCTTCT, which included the *XbaI* restriction site) and 1977 (GGCCAGTGCCAAGCTTTCACTGGATTTTGGTTTTAGGAATTAGA, which included the *HindIII* restriction site). The *mCherry-ER-Tnos* plasmid clone (Nelson, Cai and Nebenführ, 2007) was digested with *XbaI* and *HindIII* restriction enzymes to release the original *mCherry-ER-Tnos* sequence and then into the gel-purified backbone, the above-mentioned PCR fragments were cloned into it using the In-Fusion HD Cloning Plus. This effort yielded constructs harboring *LRE-cYFP-T35S* fragment or its derivatives downstream of the *35S* promoter in the *ER-r* plasmid.

### Transformation and screening of transformants

Transformations were performed as described by (Liu et al. 2016). The *pLRE::LRE-cYFP* constructs with Δ3ω and Δ5ω mutations were introduced into *lre-7* homozygous mutants. Bright, stable expressers with single insertions were identified as previously described and crossed to *ms1* to obtain *pLRE::LRE-cYFP*, *lre-7*, *ms1/+* plants. Other constructs were transformed into *Col-0*. *p35S::LRE-cYFP (WT) and p35S::LRE-cYFP (TM)* were also transformed into the *gpi8-1* mutant. Primary transformants were selected on Hygromycin (20 μg/mL; PhytoTechnology H397) for *pLRE::LRE-cYFP* and *pRBSC1A::LRE-cYFP*, or Basta (10 μg/mL; Fisher 50-240-693) for *35S* lines. YFP screening was performed in synergid cells (for *pLRE::LRE-cYFP*), root tips (for all the *35S* lines), or leaves (*pRBSC1A::LRE-cYFP* lines). At least five lines with strong fluorescence were tested for transgene expression by western blot in 9- or 14-day-old seedlings. Lines with verified protein expression and correct construct sequence were selected for biochemical assays.

### Confocal Imaging of ovules

Fluorescence imaging was performed using a Leica SP5 confocal laser scanning microscope. For cYFP detection, samples were excited with a 488 nm laser, and fluorescence emission was collected between 510 and 550 nm. Image processing and signal quantification were carried out using ImageJ software (http://imagej.nih.gov/ij/) and as described (Liu et al. 2016).

### Pollen Tube Reception Assays

*In vivo* assays to assess pollen tube reception were performed as described (Liu et al. 2016). For aniline blue staining, the protocol outlined by (Mori et al. 2006) was followed. Stained pistils were mounted in 15% glycerol, and pollen tube behavior was examined using fluorescence microscopy on a Zeiss Axiovert 100 system.

### Seed Set Assays

*In vivo* assays to assess seed set in siliques of transgenic lines carrying *LRE-cYFPΔ3ω* and *LRE-cYFPΔ5ω* were performed as described (Tsukamoto et al. 2010).

### Transient expression of proteins in *N. benthamiana*

For transient expression in *Nicotiana benthamiana*, *Agrobacterium tumefaciens* GV3101::pMP90 was electroporated with plasmids and grown overnight at 28°C in LB with appropriate antibiotics. Cultures were pelleted (5,000 rpm, 5 min), resuspended in 10 mM MgCl₂, pelleted again, and resuspended in induction buffer (100 mM MgCl₂, 10 mM MES pH 5.6, 1.5 mM acetosyringone). Bacterial suspensions were incubated at room temperature for 1–2 h to induce virulence genes before infiltration.

After incubation, bacterial suspensions were diluted to OD₆₀₀ = 0.6 in induction buffer and infiltrated into the abaxial side of fully expanded leaves of 4–6-week-old *N. benthamiana* using a needleless syringe. Plants were maintained at 21°C, 50–60% humidity, under a 16 h light / 8 h dark cycle for 9 days before imaging.

### Confocal Microscopy

Fluorescence imaging of ovules was performed as described in Liu et al. 2016. Ovule samples were imaged using a Leica SP5 confocal laser scanning microscope. For detection of cYFP, samples were excited with a 488 nm laser, and fluorescence emission was collected between 510 and 550 nm. Image acquisition and quantification were carried out using ImageJ software (https://imagej.nih.gov/ij/). After incubation, bacterial suspensions were diluted to OD₆₀₀ = 0.6 in induction buffer and infiltrated into the abaxial side of fully expanded leaves of 4–6-week-old *N. benthamiana* using a needleless syringe. Plants were maintained at 21°C, 50–60% humidity, under a 16 h light / 8 h dark cycle for 9 days before imaging.

Nine days post-infiltration, leaf sections were excised and mounted in 10% glycerol with the abaxial side facing up. Imaging was performed on a Leica SP5 confocal microscope. YFP was excited at 488 nm and detected at 510–570 nm; mCherry was excited at 543 nm and detected at 600–660 nm. Images were processed using Leica LAS AF and ImageJ software.

### Leaf Tissue collection

Tissues used for protein isolation were immediately frozen in liquid nitrogen (LN₂) and stored at −80°C. For *pRBSC1A::LRE-cYFP* and *35S* lines, 200 mg of 9-day-old or 14-day-old seedlings grown on MS plates were collected. For *gpi8-1* and the control plants, leaves were harvested from mature plants due to the slower-than-normal growth of *gpi8-1*. In the case of *pLRE::LRE-cYFP*, *lre-7*, and *ms1/+*, 100-200 mature pistils were collected after removing sepals, petals, anthers, and pedicels from the flowers.

### Protein fractionation for TP, MM, and DRM

Protein fractionation was performed as described (Sedbrook, 2002) with some modifications. Frozen tissue was ground in liquid nitrogen and homogenized in extraction buffer (Buffer 1: 50 mM potassium phosphate, pH 7.5, 2 mM MgCl₂, 50 mM NaCl, and 1× protease inhibitor cocktail [Millipore, 539131]) at a buffer:tissue ratio of ≥3:1 for seedlings and 1:1 for pistils (200 pistils/genotype). After centrifugation at 10,000 g, 4°C for 10 min, the supernatant was collected as total protein (TP). For microsomal membrane (MM) isolation, the TP was centrifuged at 100,000 g, 4°C for 3 h. The resulting pellet was washed and resuspended in TBS. The pellet was then washed once with 150 μL Tris-buffered saline (TBS, 10 mM Tris HCl, pH 7.5, 150 mM NaCl, referred to as buffer 2 in Supplementary Fig. S3 (Doering, Englund and Hart, 1993) and resuspended with the same TBS buffer to yield the MM.

To isolate detergent-resistant membranes (DRMs), ¼ volume of precondensed Triton X-114 (Doering, Englund and Hart, 1993) was added to MM, incubated on ice for 15 min, and centrifuged at 10,000 g, 4°C for 10 min. The supernatant was incubated at 37°C for 10 min and centrifuged at 1,000 g (33–37°C) for 10 min. The lower phase (LP) was collected, adjusted to original volume with TBS, and subjected to two additional rounds of phase separation. The final lower phase (LP3) was designated as the DRM fraction.

### Western blotting

Protein concentrations of total (TP) and microsomal membrane (MM) fractions were determined using the Bradford Protein Assay (Bio-Rad, #5000006) with a BioTek™ Eon™ microplate spectrophotometer and Gen5™ v2.0 software. Unless otherwise specified, 10 µg of TP and MM protein were loaded per lane. Due to interference from detergents and hydrophobic content, DRM (LP3) protein concentration could not be quantified by Bradford; instead, 10 µL of a 1:10 dilution of LP3 was loaded. For gpi8-1 samples, protein loading was normalized based on western blot signals using an anti-GFP antibody (Covance MMS-118P).

Protein was separated in 12% gel (Acrylamide/Bis, J.T.Baker, 4970-00) with 120 V voltage for 2.5 h-3 h, and transferred (0.3 A, 1.5 h) to PVDF (Immobilon-P Membrane, emd millipore, IPVH10100). The blot was stained by 0.1% (w/v) Ponceau S (in 5% acetic acid) for about 5 minutes, imaged and then washed several times with 1× PBST (100 mM NaCl, 80 mM Na2HPO4, and 20 mM NaH2PO4, pH 7.5, 0.1% (w/v) detergent of Tween-20) before blocking with 5% nonfat milk in 1× PBST buffer for one hour. The antibody incubation was done using anti-GFP (1:1000 for 1-1.5 hours) and anti-mouse (1:7500 for 1-1.5 hours, goat anti-mouse IgG-HRP, Santa Cruz Biotechnology, sc-2005) followed by three times washing (1× PBST, 5min/time) after the primary antibody incubation. Alternatively, anti-SKU5 (1:1000 for one hour) and goat anti-rabbit (1:7500 for one hour, goat anti-rabbit IgG-HRP, Santa Cruz Biotechnology, sc-2004) were used. For re-probing, a mild stripping buffer was used following the mild stripping protocol from abcam (http://www.abcam.com/). UGPase UDP-glucose pyrophosphorylase antibody was from Agrisera, Q43772. ECL (Amersham ECL Select Western blotting detection reagent, RPN2235) was used to develop the membrane. The final images were taken using Azure c300 (Azure Biosystems).

### PI-PLC treatment

PI-PLC treatment was performed as described (Sedbrook et al. 2002) with minor modifications. DRM fractions were diluted 1:10 and incubated with 1 unit/100 µL phosphatidylinositol-specific phospholipase C (PI-PLC; Sigma P5542) at 30 °C for 1.5 h with intermittent mixing. Post-incubation, samples were centrifuged at 1,000 g above 33 °C for 10 min. The upper phase (UP) was collected and subjected to a second Triton X-114 extraction (Doering, Englund and Hart, 1993).

Proteins in the final UP were precipitated using 10% trichloroacetic acid (Alfa Aesar, 22156) for 2 h on ice, followed by centrifugation at maximum speed for 10 min at 4 °C. Bovine serum albumin (1 µg) was added as a recovery control prior to precipitation. Pellets were resuspended in SDS loading buffer, and pH was neutralized using ammonium hydroxide (VWR 470300-210) until the bromophenol blue indicator turned blue. Three biological replicates were performed per construct from tissue harvest through PI-PLC treatment.

---

**Supplementary Fig. S1.** Transgenic constructs used in this study for expression in the synergid cells of the female gametophyte.

A. Diagram of the LRE-cYFP protein. Wild-type or altered amino acid sequence of the predicted ω amino acid in each construct is indicated below the diagram. Each dash in the protein sequence represents a deletion of the corresponding amino acid in wild-type LRE protein sequence. SP, signal peptide, GAS, GPI attachment signal (lighter gray rectangle).

B. Diagrams depicting the constructs utilized to express the full length (LRE-cYFP) or the ω sites deletions–3ω or 5ω from the *LRE* promoter. The length of each portion in the diagram is proportional to its actual length in the protein. The wild-type *LRE* sequence is used in the *pLRE::LRE-cYFP*. In the *pLRE:LRE-cYFPΔ3ω*, the top 3 predicted ω sites by the metazoan Pi predictor were deleted. In the *pLRE:LRE-cYFPΔ5ω* construct, the top 3 ω sites predicted by both prediction sites were removed. ω, omega site; GAS, GPI attachment signal.

**Supplementary Fig. S2.** Deletion of predicted ω-sites in LRE does not impair pollen tube reception or fertility.

A – C. Representative images showing pollen tube reception in ovules of indicated genotype when crossed to *pLAT52:GUS* pollen. This GUS staining based assay was used to score pollen tube reception. Bar = 50 μm.

D. The *pLRE:LRE-cYFPΔ3ω* and *pLRE:LRE-cYFPΔ5ω* constructs fully complement the pollen tube reception defect in *lre-7/lre-7* plants. Total number of ovules analyzed are in the center of each column. Number below each column, two independent transformants (*pLRE:LRE-cYFPΔ3ω*) and three independent transformants (*pLRE:LRE-cYFPΔ5ω*) that are single insertion and homozygous for the respective transgenes in the *lre-7/lre-7* background.

E. Light micrographs of wild-type and *lre/lre* siliques, opened at 10 days after pollination (DAP). In lre/lre siliques, normal seeds (#), undeveloped ovules (black arrow) and aborted seeds (asterisk) are marked. Scale bars, 500μm.

F. The *pLRE:LRE-cYFPΔ3ω* and *pLRE:LRE-cYFPΔ5ω* constructs fully complement the seed set defects in *lre-7/lre-7* plants. Total number of seeds analyzed are in the center of each column. Number below each column, two independent transformants (*pLRE:LRE-cYFPΔ3ω*) and three independent transformants (*pLRE:LRE-cYFPΔ5ω*) that are single insertion and homozygous for the respective transgenes in the *lre-7/lre-7* background.

**Supplementary Fig. S3.** Transgenic constructs used in this study for ectopic expression of LRE-cYFP in seedlings.

A. Diagram of the LRE-cYFP protein. Wild-type or altered amino acid sequence of the predicted ω amino acid in each construct is indicated below the diagram. Each dash in the protein sequence represents a deletion of the corresponding amino acid in wild-type LRE protein sequence. In LRE-cYFP-TM, predicted single transmembrane region of FERONIA is shown in gray.

B. The wild-type *LRE* sequence is used in *pRBSC1A::LRE-cYFP* and *p35S::LRE-cYFP* constructs. In the *p35S::LRE-cYFPΔ2ω* construct, the ω site and the second-best predicted ω site are deleted. In the *p35S::LRE-cYFPΔGAS* construct, the GAS domain is removed. In the *p35S::LRE-cYFP-TM* construct, both ω site and GAS domain are replaced with FERONIA’s transmembrane domain (TM), a single transmembrane domain-containing receptor-like kinase.

**Supplementary Fig. S4.** Flowchart illustrating the process from tissue collection to PI-PLC treatment.

Protein subjected to PI-PLC treatment was prepared in four steps. In Step 1, total protein (TP) was extracted. Tissue (9- or 14-day-old seedlings without roots) was ground into a fine powder in liquid nitrogen, homogenized in an extraction buffer (Buffer 1), and centrifuged to remove insoluble debris (green pellet). The supernatant, referred to as TP, was collected. In Step 2, microsomal membrane (MM) protein was isolated by high-speed centrifugation, washed once with Tris-buffered saline (TBS, Buffer 2), and resuspended in the same buffer. The supernatant (SUP) from this centrifugation was also collected. In Step 3, Triton X-114 (TX-114) treatment was performed thrice to isolate detergent-resistant membranes (DRM). After each detergent treatment and centrifugation, the upper phase (UP) and lower phase (LP) were separately collected. The third LP fraction (LP3) was designated as DRM. In Step 4, DRM was diluted tenfold and split into two samples: one treated with PI-PLC and the other with a mock buffer as a control. After treatment and centrifugation, a new set of upper and lower phases was generated (UP4 and LP4). **“+“** indicates treatment with PI-PLC, while **“–“** indicates the mock-treated control.

TP, total protein; SUP, supernatant; MM, microsomal membrane; TX-114, precondensed Triton X-114; UP, upper phase; LP, lower phase; DRM, detergent-resistant membrane.

**Supplementary Fig. S5.** Expression of *LRE-cYFP* and variants in Arabidopsis Col-0 leaves under the *35S* promoter.

Expression of *AtLRE*-cYFP, *AtLRE*-cYFP-TM, *AtLRE*-cYFPΔ2ω, and *AtLRE*-cYFPΔGAS in Arabidopsis *Col-0* leaves (14–16 days old). Imaging was performed using bright-field (BF) and GFP fluorescence (GFP) channels.

**Supplementary Fig. S6.** Western blotting of TP, MM, and SUP protein fractions from different LRE variants.

A. LRE total protein of different variants from 9 or 14 day-old seedlings. The non-transgenic (NT) (*Col*-0) seedling protein was used as a negative control. The bracket shows bands of LRE-cYFP variants.

B. LRE protein in MM and SUP from LRE-cYFP (WT) and LRE-cYFPΔGAS (ΔGAS) variants. Total protein from the LRE-cYFP construct was used as an internal control. Step # refers to the steps in Supplementary Fig. S2.

ΔGAS, LRE-cYFPΔGAS, LRE protein without predicted GPI-attachment signal (GAS); WT, LRE-cYFP; TP, total protein; MM, microsomal membrane proteins; SUP, supernatant.

