## Supplemental Data combined for "Structural Determinants and Biochemical Characterization of LORELEI as a GPI-Anchored Protein"

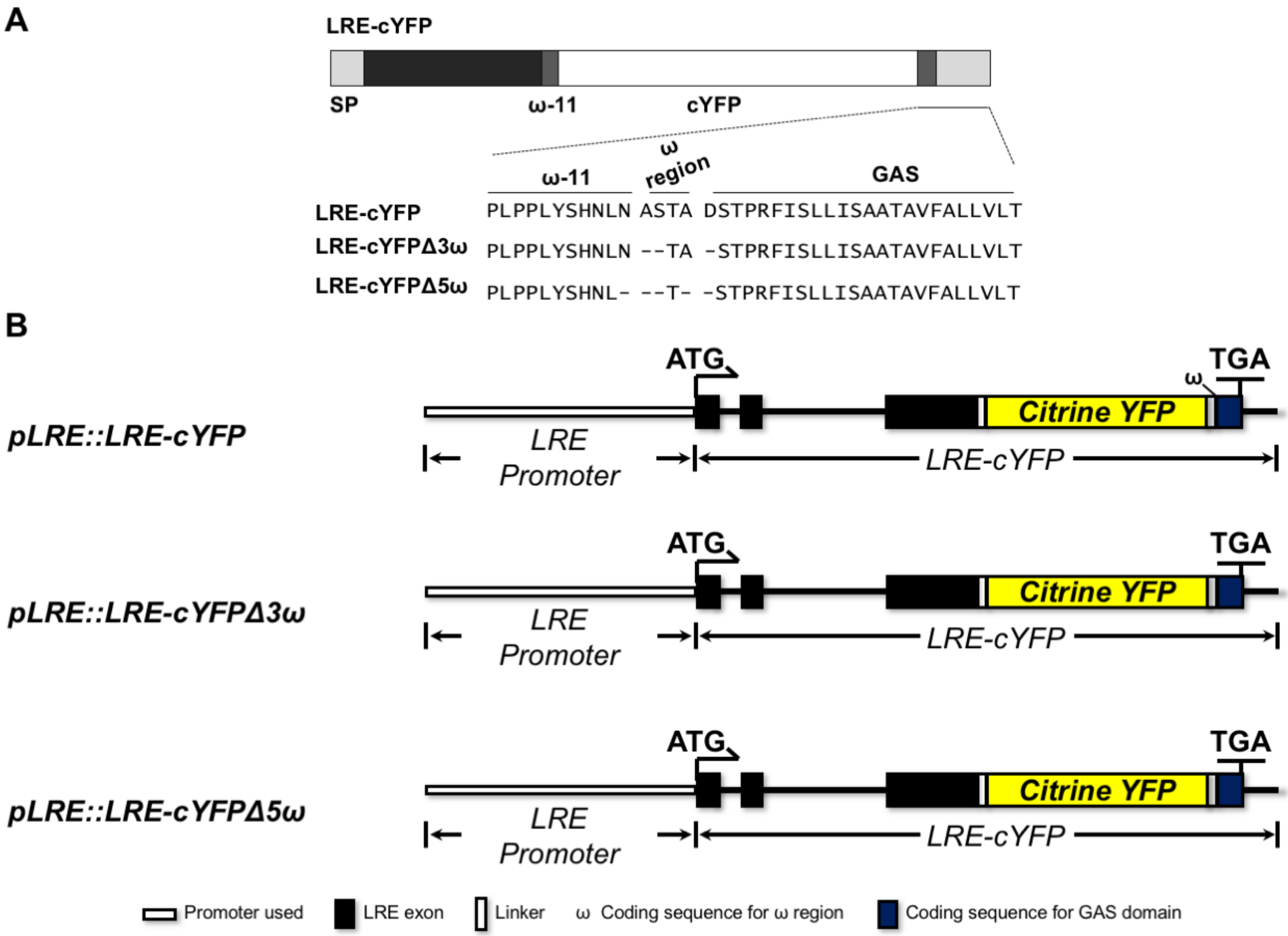
A
LRE-cYFP
SP
ω-11
cYFP
ω-11
ω region
GAS
LRE-cYFP PLPPLYSHNLN ASTA DSTPRFISLLISAATAVFALLVLT
LRE-cYFPΔ3ω PLPPLYSHNLN --TA -STPRFISLLISAATAVFALLVLT
LRE-cYFPΔ5ω PLPPLYSHNL- --T- -STPRFISLLISAATAVFALLVLT
B
pLRE::LRE-cYFP
pLRE::LRE-cYFPΔ3ω
pLRE::LRE-cYFPΔ5ω
ATG
TGA
ω
Citrine YFP
LRE Promoter
LRE-cYFP
Promoter used
LRE exon
Linker
ω Coding sequence for ω region
Coding sequence for GAS domain

Fig. S1

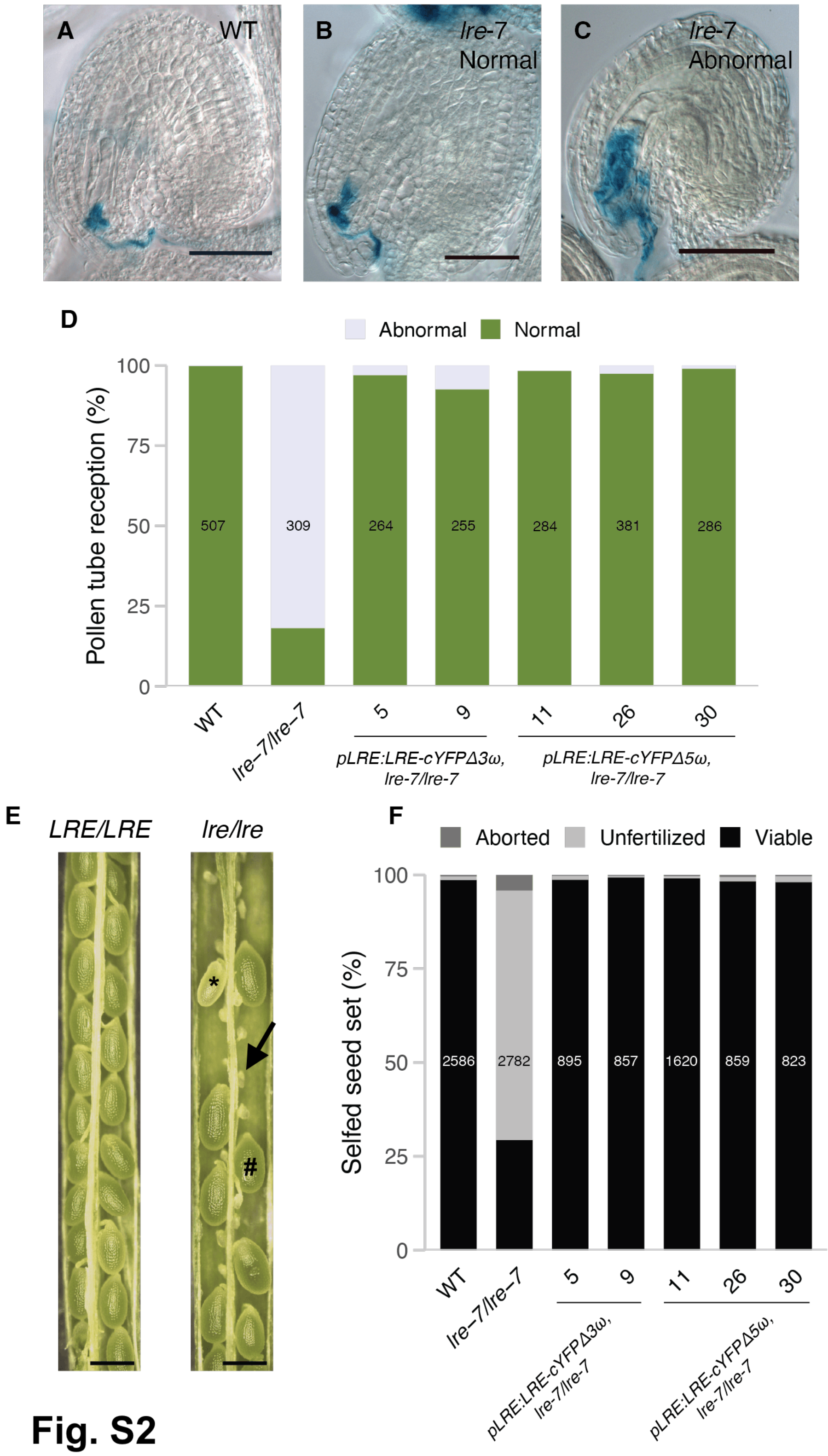

A
WT
B
lre-7
Normal
C
lre-7
Abnormal
D
Abnormal
Normal
Pollen tube reception (%)
100
75
50
25
0
507
309
264
255
284
381
286
WT
lre–7/lre–7
5
9
11
26
30
pLRE:LRE-cYFPΔ3ω, lre-7/lre-7
pLRE:LRE-cYFPΔ5ω, lre-7/lre-7
E
LRE/LRE
lre/lre
*
#
F
Aborted
Unfertilized
Viable
Selfed seed set (%)
100
75
50
25
0
2586
2782
895
857
1620
859
823
WT
lre–7/lre–7
5
9
11
26
30
pLRE:LRE-cYFPΔ3ω, lre-7/lre-7
pLRE:LRE-cYFPΔ5ω, lre-7/lre-7


**Fig. S2**

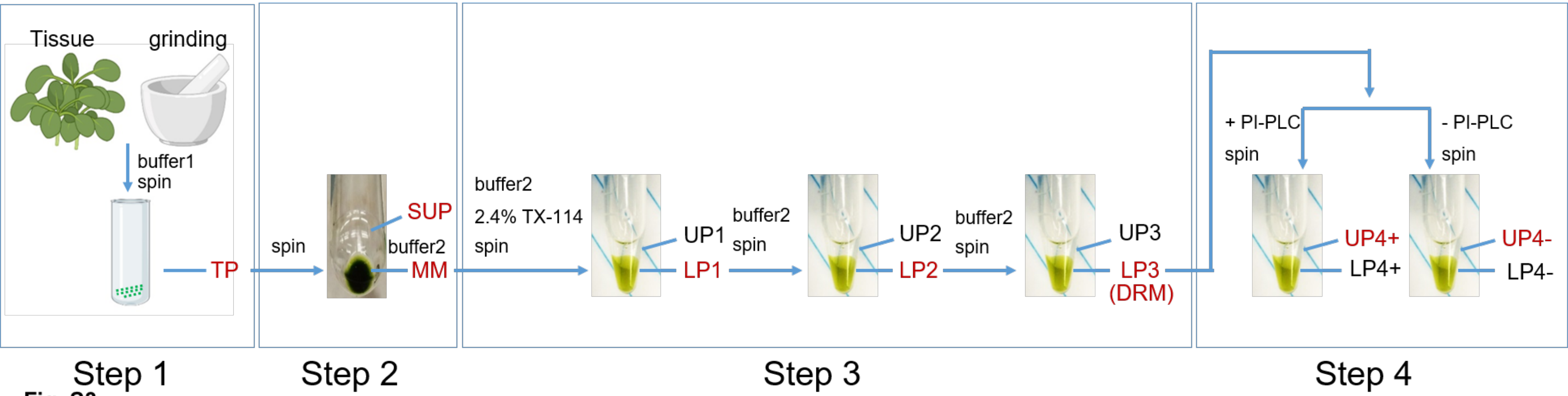
Tissue
grinding
buffer1
spin
TP
spin
SUP
buffer2
MM
buffer2
2.4% TX-114
spin
UP1
LP1
buffer2
spin
UP2
LP2
buffer2
spin
UP3
LP3
(DRM)
+ PI-PLC
spin
- PI-PLC
spin
UP4+
LP4+
UP4-
LP4-
Step 1
Step 2
Step 3
Step 4

**Fig. S3**

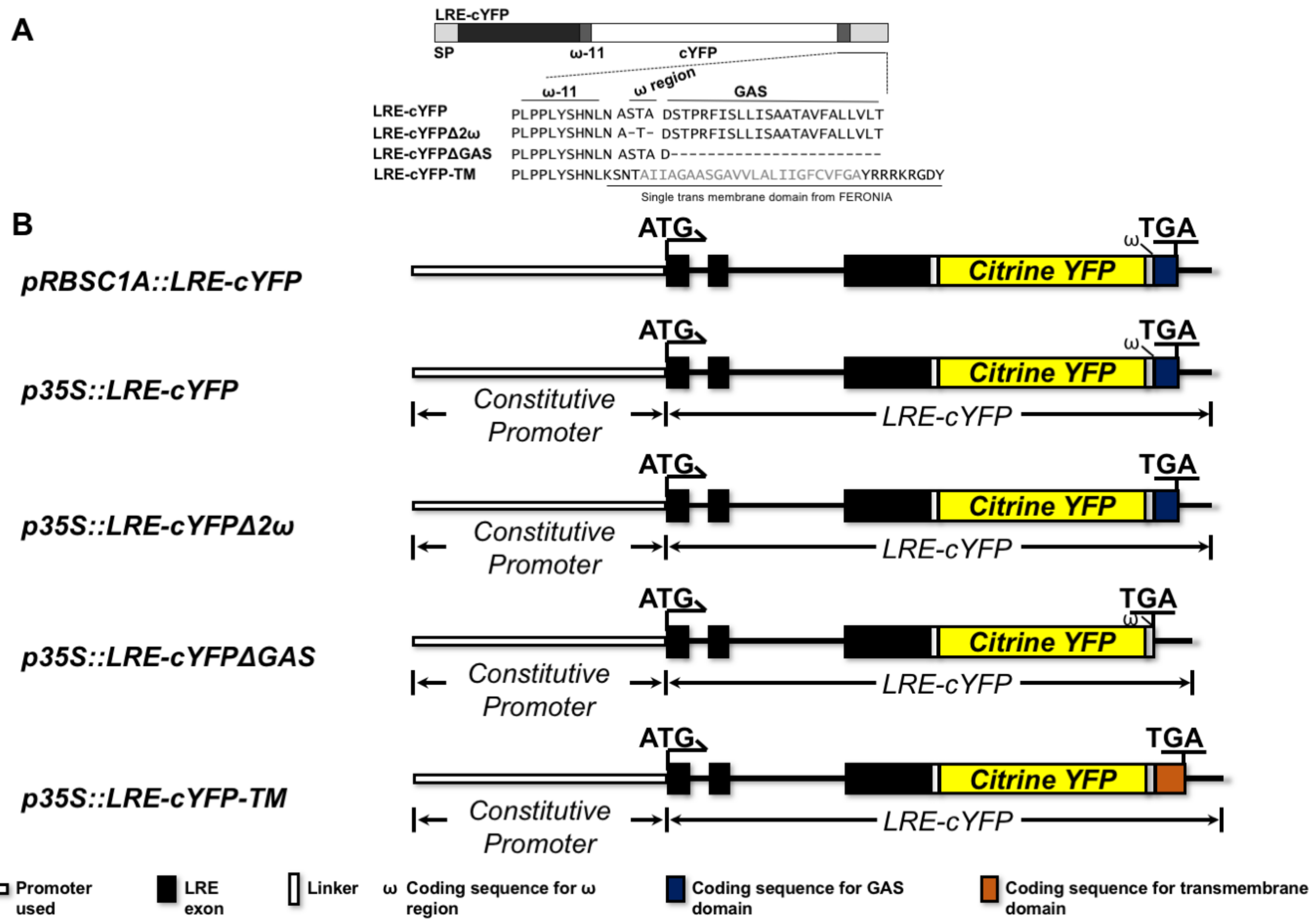
A
LRE-cYFP
SP
ω-11
cYFP
ω-11
ω region
GAS
LRE-cYFP PLPPLYSHNLN ASTA DSTPRFISLLISAATAVFALLVLT
LRE-cYFPΔ2ω PLPPLYSHNLN A-T- DSTPRFISLLISAATAVFALLVLT
LRE-cYFPΔGAS PLPPLYSHNLN ASTA D-----------------------
LRE-cYFP-TM PLPPLYSHNLKSNTAIIAGAASGAVVLALIIGFCVFGAYRRRKRGDY
Single trans membrane domain from FERONIA
B
pRBSC1A::LRE-cYFP
p35S::LRE-cYFP
p35S::LRE-cYFPΔ2ω
p35S::LRE-cYFPΔGAS
p35S::LRE-cYFP-TM
ATG
TGA
ω
Citrine YFP
Constitutive Promoter
LRE-cYFP
Promoter used
LRE exon
Linker
ω Coding sequence for ω region
Coding sequence for GAS domain
Coding sequence for transmembrane domain

**Fig. S4**

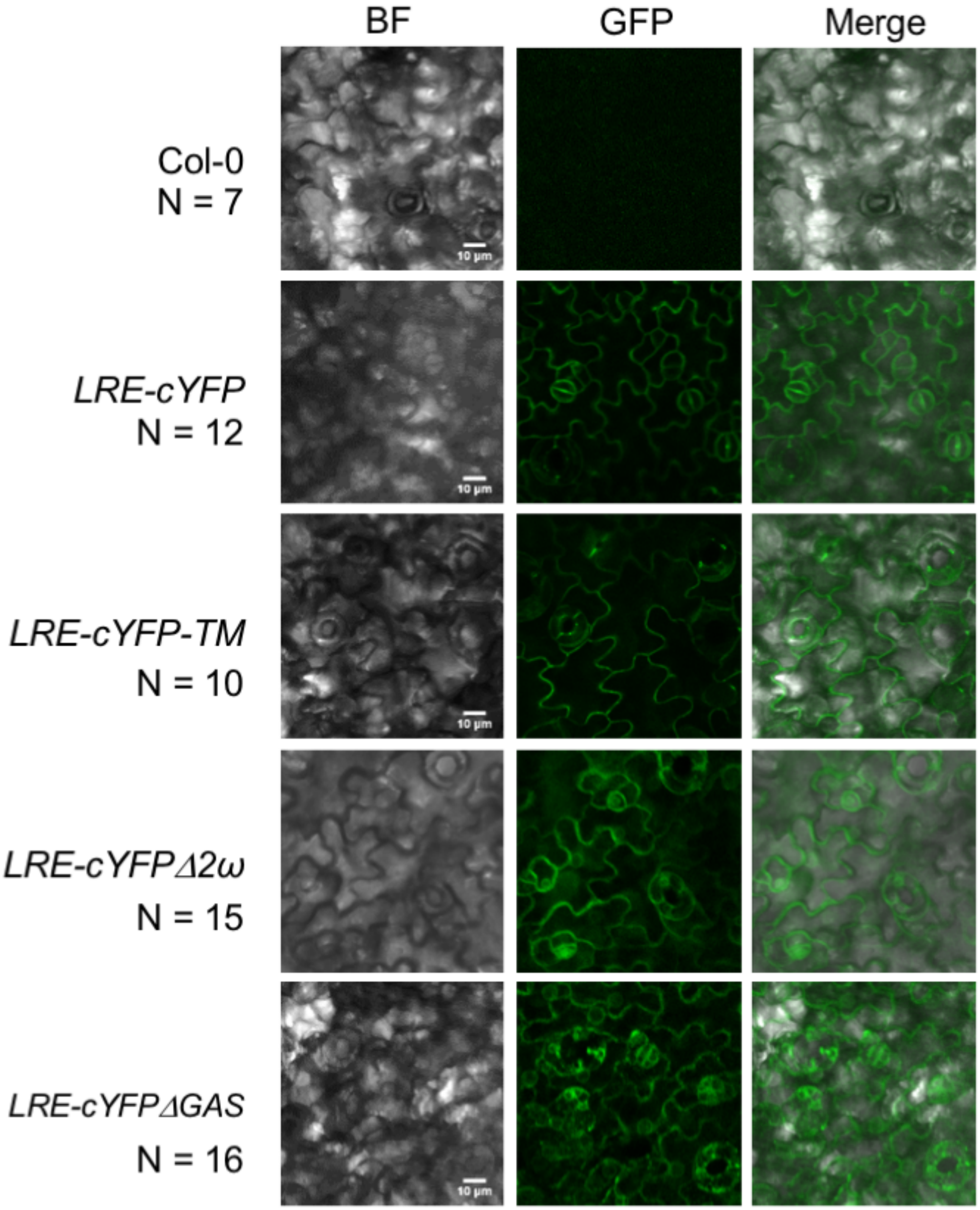
BF
GFP
Merge
Col-0
N = 7
LRE-cYFP
N = 12
LRE-cYFP-TM
N = 10
LRE-cYFPΔ2ω
N = 15
LRE-cYFPΔGAS
N = 16

**Fig. S5**

Fig. S6

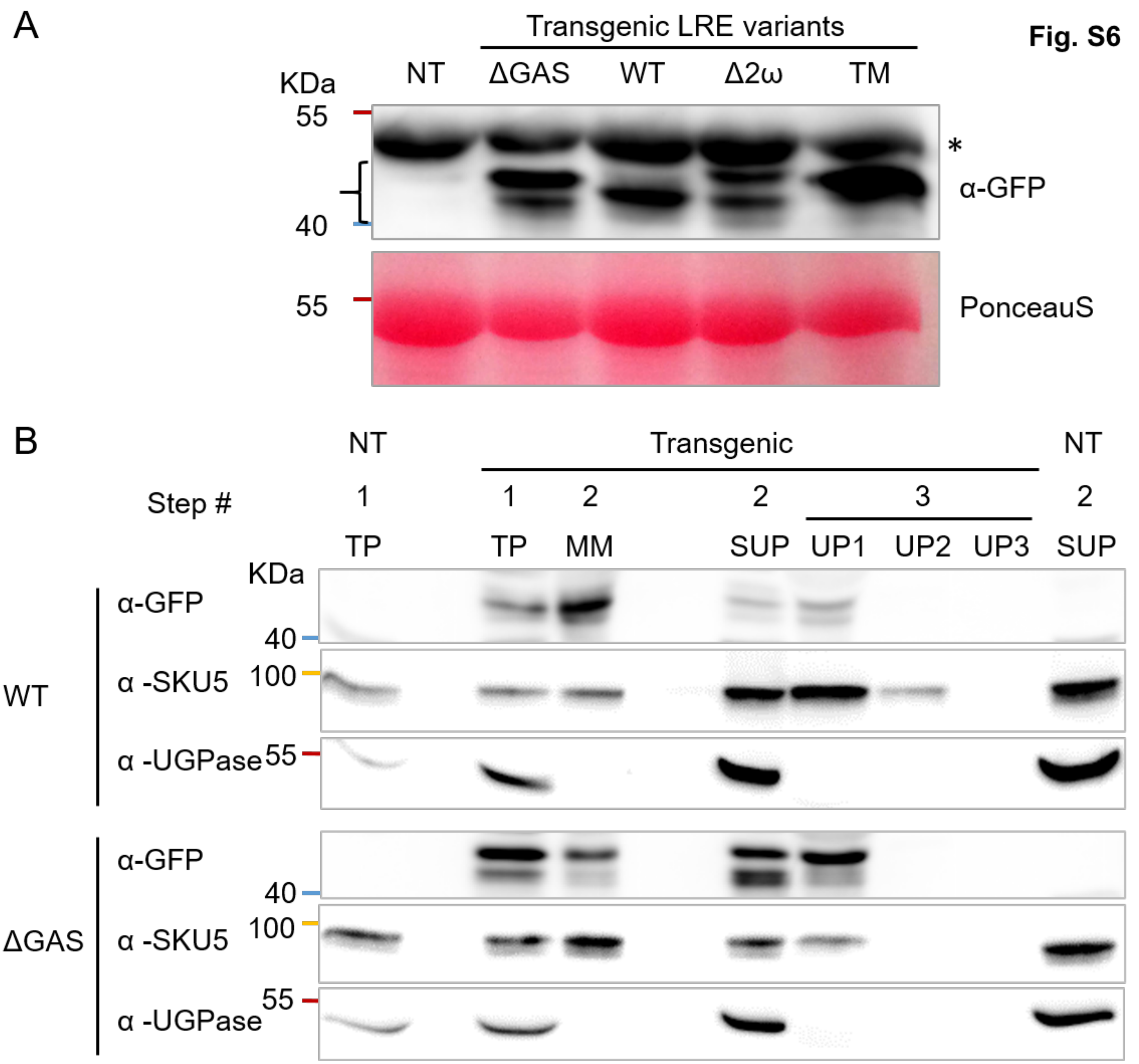
A
Transgenic LRE variants
KDa
NT
ΔGAS
WT
Δ2ω
TM
55
40
*
α-GFP
55
PonceauS
B
NT
Transgenic
NT
Step #
1
1
2
2
3
2
TP
TP
MM
SUP
UP1
UP2
UP3
SUP
KDa
WT
α-GFP
40
α -SKU5
100
α -UGPase
55
ΔGAS
α-GFP
40
α -SKU5
100
α -UGPase
55

**Supplementary Table S2. Predicted protein products, sizes, and expected localization of LRE variants used in this study.**

| Background genotype | LRE-cYFP variant | Protein type | # of amino acids[1] | Predicted size of protein (kDa) (@.110 kDa / amino acid)[2] | Predicted localization after biochemical fractionation |
|---|---|---|---|---|---|
| *Columbia Wildtype* | ΔGAS | LRE- cYFP | 382 | 42.02 | Non-MM |
| | WT | LRE-cYFP-GAS | 405 | 44.55 | MM |
| | | LRE-cYFP-GPI | 378 + GPI* | 42.69-44.69 | MM |
| | | LRE-cYFP | 379 | 41.69 | Non-MM |
| | Δ2ω | LRE-cYFP-GAS | 403 | 44.33 | MM |
| | | LRE-cYFP-GPI | 380 + GPI* | 42.80-44.80 | MM |
| | | LRE-cYFP | 380 | 41.80 | Non-MM |
| | TM | LRE-cYFP-TM | 413 | 45.43 | MM |
| *gpi8-1* | WT | LRE-cYFP-GAS | 405 | 44.55 | MM |
| | | LRE-cYFP-GPI | 379 + GPI* | 42.69-44.69 | MM |
| | | LRE-cYFP | 379 | 41.69 | Non-MM |
| | TM | LRE-cYFP-TM | 413 | 45.43 | MM |

[1]Estimated size of the pre-protein in the ER after the N-terminal signal sequence is processed, that is removed, but includes the GAS sequence.

[2]Estimated molecular weight of a plant amino acid (https://www.thermofisher.com/us/en/home/references/ambion-tech-support/rna-tools-and-calculators/proteins-and-amino-acids.html).

WT, wild type LRE sequence; GAS, GPI attachment signal sequence; Δ2ω, deletion of top two predicted ω sites in LRE (Figure 1A, 1B, Supplementary Figure 1); MM, microsomal membrane fraction (Supplementary Figure S3); TM, a single transmembrane domain of FERONIA (Supplementary Figure 1).

*Estimated molecular weight of a plant GPI is 1-3 kDa.

**Supplementary Table S1.** Enhanced transmission of transgene either with Δ3ω or Δ5ω mutations through the *lre-7* female gametophyte indicates complementation of the *lre-7* reproductive defects by the introduced transgene.

| Female parent⁺ | Male parent⁺ | Observed No. of progeny | | TE (R/S) | χ2† | P-value |
|---|---|---|---|---|---|---|
| | | $Hyg^{R*}$ | $Hyg^{S*}$ | | | |
| *pLRE:LRE-cYFPΔ3ω/+*, *lre-7/lre-7* | | | | | | |
| Col-0 | Line 5 | 171 | 204 | 0.838 | 2.90 | 0.088# |
| Line 5 | Col-0 | 222 | 14 | 15.86 | 183.32 | < 0.001 |
| Col-0 | Line 9 | 204 | 194 | 1.05 | 0.251 | 0.616# |
| Line 9 | Col-0 | 226 | 22 | 10.27 | 167.81 | <0.001 |
| *pLRE:LRE-cYFPΔ5ω/+*, *lre-7/lre-7* | | | | | | |
| Col-0 | Line 11 | 159 | 203 | 0.78 | 5.35 | 0.020 |
| Line 11 | Col-0 | 193 | 18 | 10.722 | 145.14 | <0.001 |
| Col-0 | Line 26 | 133 | 178 | 0.747 | 6.51 | 0.010 |
| Line 26 | Col-0 | 231 | 18 | 12.83 | 182.21 | <0.001 |
| Col-0 | Line 30 | 217 | 202 | 1.07 | 0.54 | 0.464# |
| Line 30 | Col-0 | 194 | 9 | 21.55 | 168.60 | <0.001 |

Col-0, non-transgenic, Columbia wild-type plants.

⁺ Line numbers refer to independent transformants in the *lre-7/lre-7* background containing single insertion of indicated mutant transgene. Genotype of each transgenic line used is heterozygous for the indicated mutant transgene (*mutant transgene/+*) and homozygous for the *lre-7* mutation (*lre-7/lre-7*).

*Hygromycin resistant ($Hyg^R$) and susceptible ($Hyg^S$) progeny. Hygromycin resistance gene is linked to the construct carrying the indicated transgene.

TE, Transmission efficiency was calculated as the ratio of hygromycin resistance (R) to susceptibility (S) in the progeny of the indicated cross.

†χ2 is calculated based on an expected segregation ratio of hygromycin resistant to susceptibility of 1:1.

#No significant deviation from 1:1 segregation through the male gametophyte indicates that pollen parent contains a single insertion of the indicated mutant transgene. Additional details on the protocol used to isolate single insertion lines can be found in Liu *et al.,* 2016.
